# Multiple levels of transcriptional regulation control glycolate metabolism in *Paracoccus denitrificans*

**DOI:** 10.1101/2024.03.11.584432

**Authors:** Lennart Schada von Borzyskowski, Lucas Hermann, Katharina Kremer, Sebastian Barthel, Bianca Pommerenke, Timo Glatter, Nicole Paczia, Erhard Bremer, Tobias J. Erb

## Abstract

The hydroxyacid glycolate is a highly abundant carbon source in the environment. Glycolate is produced by unicellular photosynthetic organisms and excreted at petagram scales to the environment, where it serves as growth substrate for heterotrophic bacteria. In microbial metabolism, glycolate is first oxidized to glyoxylate by the enzyme glycolate oxidase. The recently described β-hydroxyaspartate cycle (BHAC) subsequently mediates the carbon-neutral assimilation of glyoxylate into central metabolism in ubiquitous Alpha- and Gammaproteobacteria. While the reaction sequence of the BHAC was elucidated in *Paracoccus denitrificans*, little is known about the regulation of glycolate and glyoxylate assimilation in this relevant alphaproteobacterial model organism. Here, we show that regulation of glycolate metabolism in *P. denitrificans* is surprisingly complex, involving two regulators, the IclR-type transcription factor BhcR that acts as an activator for the BHAC gene cluster, as well as the GntR-type transcriptional regulator GlcR, a previously unidentified repressor that controls the production of glycolate oxidase. Furthermore, an additional layer of regulation is exerted at the global level, which involves the transcriptional regulator CceR that controls the switch between glycolysis and gluconeogenesis in *P. denitrificans*. Together, these regulators control glycolate metabolism in *P. denitrificans*, allowing the organism to assimilate glycolate together with other carbon substrates in a simultaneous fashion, rather than sequentially. Our results show that the metabolic network of Alphaproteobacteria shows a high degree of flexibility to react to the availability of multiple substrates in the environment.

## Introduction

The two-carbon compound glycolate is the simplest α-hydroxyacid. Photosynthetic organisms that rely on carbon dioxide fixation via the Calvin-Benson-Bassham (CBB) cycle produce this molecule as part of their photorespiration process (1). Subsequently, glycolate can either be recycled into cellular metabolism using an inefficient and energetically costly metabolic pathway (2) or excreted (3). The latter route predominates in unicellular photosynthetic organisms, such as eukaryotic microalgae and Cyanobacteria. Due to the abundance of these ubiquitous phototrophs in marine and freshwater habitats, an annual flux of one petagram (10^9^ tonnes) of glycolate has been estimated (4). Hence, glycolate is a readily available source of carbon for heterotrophic environmental microorganisms.

In microbial metabolism, glycolate is first oxidized to glyoxylate via the enzyme glycolate oxidase (5-7). In addition, glyoxylate can be generated as breakdown product of ubiquitous purine bases and allantoin (8), as well as ethylenediaminetetraacetate (EDTA) and nitrilotriacetate (NTA) (9, 10). There are two known metabolic routes for subsequent net assimilation of glyoxylate. The well-studied glycerate pathway is used by *Escherichia coli* and other bacteria to convert two molecules of glyoxylate into one molecule of 2-phosphoglycerate, releasing one molecule of carbon dioxide in the process (7, 11). The β-hydroxyaspartate cycle (BHAC) (12, 13), on the other hand, was recently shown to convert two molecules of glyoxylate into oxaloacetate via four enzymatic steps without the release of CO_2_ (14). In contrast to the glycerate pathway, the BHAC has a much higher energy and carbon efficiency, and has already been successfully applied in metabolic engineering efforts in bacteria (15) and plants (16).

Notably, the BHAC is the dominant glycolate assimilation route in the environmentally relevant groups of Alpha- and Gammaproteobacteria, and was recently shown to play an important role in global glycolate conversions, in particular in marine environments (14). In a field study, enzymes of the BHAC were shown to be upregulated during a bloom of marine algae, following increased glycolate concentrations. Metagenomic data further supported the global prevalence of the BHAC. However, despite the ecological relevance of the BHAC, the question how glycolate and glyoxylate metabolism are regulated at the molecular and cellular level in Alpha- and Gammaproteobacteria remained unanswered.

The BHAC was previously elucidated in the Alphaproteobacterium *Paracoccus denitrificans* (14), an environmental model organism with a versatile metabolism (17, 18). *P. denitrificans* can grow both aerobically and anaerobically, using either oxygen or nitrate as terminal electron acceptor (19, 20). In addition, *P. denitrificans* is capable of utilizing many different carbon substrates for heterotrophic growth and can even fix carbon dioxide for autotrophic growth (21, 22). While the regulation of denitrification (23-26) and respiration (27, 28) were elucidated in detail in *P. denitrificans*, the mechanisms that regulate central carbon metabolism in this bacterium have been studied only recently (29-32).

In respect to glycolate metabolism, it is known that production of the four enzymes of the BHAC is strongly induced in *P. denitrificans* during growth on glycolate, compared to growth on succinate. Furthermore, it was reported that *bhcR*, a gene coding for a putative transcriptional regulator, is positioned adjacent to the enzyme-encoding genes of the BHAC in the genome of *P. denitrificans*. BhcR was found to bind to the promoter region of the *bhc* gene cluster, while, in turn, this interaction was negatively affected by glyoxylate (14).

In this work, we show that BhcR functions as an activator of the *bhc* gene cluster and is required for both growth on glyoxylate and glycolate in *P. denitrificans*. In addition, we identify and characterize GlcR, a previously unknown transcriptional repressor of the GntR family that regulates glycolate oxidase in *P. denitrificans*. By extending our investigation to the global level, we found that the transcription factor CceR controls the metabolic switch between glycolysis and gluconeogenesis. Furthermore, we show that *P. denitrificans* co-assimilates glycolate and other carbon substrates simultaneously, not sequentially. Collectively, our work demonstrates multiple levels of transcriptional regulation in glycolate metabolism and highlights the surprising flexibility of the central metabolic network of Alphaproteobacteria in response to carbon substrate availability.

## Materials & Methods

### Chemicals & Reagents

Unless otherwise stated, all chemicals and reagents were acquired from Sigma-Aldrich (St. Louis, USA), and were of the highest purity available. Sodium glycolate was acquired from Alfa Aesar (Haverhill, USA).

### Strains, media and cultivation conditions

All strains used in this study are listed in **Supplementary Table 4**. *Escherichia coli* DH5α (for genetic work), ST18 (33) (for plasmid conjugation to *P. denitrificans*) and BL21 AI (for protein production) were grown at 37 °C in lysogeny broth (34), unless stated otherwise.

*Paracoccus denitrificans* DSM 413 (35) and its derivatives were grown at 30 °C in lysogeny broth or in mineral salt medium with TE3-Zn trace elements (36) supplemented with various carbon sources. To monitor growth, the OD_600_ of culture samples was determined on a photospectrometer (Merck Chemicals GmbH, Darmstadt, Germany) or in Infinite® 200 Pro plate reader systems (Tecan, Männedorf, Switzerland).

### Vector construction

All plasmids used in this study are listed in **Supplementary Table 5**.

To create a plasmid for heterologous overexpression of *glcR* in *E. coli*, this gene (Pden_4400) was cloned into the expression vector pET16b (Merck Chemicals). To this end, the respective gene was amplified from genomic DNA of *P. denitrificans* DSM 413 using the primers provided in **Supplementary Table 6**. The resulting PCR product was digested with suitable restriction endonucleases (Thermo Fisher Scientific) as given in **Supplementary Table 6** and ligated into the expression vector pET16b that had been digested with the same enzymes to create a vector for heterologous production of GlcR. The gene encoding for BhcR (Pden_3922) had been cloned into pET16b previously (14). To heterologously produce MBP-GlcR, the *glcR* gene was codon-optimized using Geneious Prime (Biomatters, Inc., Boston, USA) and ordered from Twist Bioscience (South San Francisco, USA), including terminal *Bsm*BI endonuclease sites. This fragment was inserted into an expression vector (pMBP-sfgfp_dropout) encoding for an N-terminal maltose-binding protein gene (*malE*) by Golden Gate assembly with the *Bsm*BI isoschizomer *Esp*3I.

To create constructs for gene deletion in *P. denitrificans*, the upstream and downstream flanking regions of the *bhcR*/*glcR*/*cceR*/*glcDEF* genes from *P. denitrificans* DSM 413 were cloned into the gene deletion vector pREDSIX (37). To this end, the flanking regions were amplified from genomic DNA of *P. denitrificans* DSM 413 with the primers given in **Supplementary Table 6**. The resulting PCR products were used to perform Gibson assembly with the vector pREDSIX, which had been digested with *Mfe*I. Subsequently, the resulting vector was digested with *Nde*I, and a kanamycin resistance cassette, which had been cut out of the vector pRGD-Kan (37) with *Nde*I, was ligated into the cut site to generate the final vectors for gene deletion. In each case, vectors with forward orientation and reverse orientation of the kanamycin resistance cassette were generated.

To create the promoter probe vector pTE714, the mCherry gene was amplified with the primers mCherry_fw and mCherry_rv using the vector pTE100-mCherry (38) as template. The PCR product was digested with *Nde*I and *Eco*RI and subsequently ligated into the backbone of pTE100 (38) (digested with *Ase*I and *Mfe*I), yielding the pTE714 plasmid.

To create reporter plasmids for *P. denitrificans*, the intergenic regions between *bhcR*/*bhcA* (Pden_3922/Pden_3921) and *glcR*/*glcD* (Pden_4400/4399), respectively, were cloned into the promoter probe vector pTE714. The respective regions were amplified from genomic DNA of *P. denitrificans* DSM413 with the primers provided in **Supplementary Table 6**. The resulting PCR products were digested with suitable restriction endonucleases (Thermo Fisher Scientific, Waltham, USA) as given in **Supplementary Table 6** and ligated into likewise digested pTE714.

Successful cloning of all desired constructs was verified by Sanger sequencing (Microsynth, Göttingen, Germany).

### Production and purification of recombinant proteins

For heterologous overproduction of BhcR and GlcR, the corresponding plasmid encoding the respective protein was first transformed into chemically competent *E. coli* BL21 AI cells. The cells were then grown on LB agar plates containing 100 µg mL^-1^ ampicillin at 37 °C overnight. A starter culture in selective LB medium was inoculated from a single colony on the next day and left to grow overnight at 37 °C in a shaking incubator. The starter culture was used on the next day to inoculate an expression culture in selective TB medium in a 1:100 dilution. The expression culture was grown at 37 °C in a shaking incubator to an OD_600_ of 0.5 to 0.7, induced with 0.5 mM IPTG and 0.2% L-arabinose and subsequently grown overnight at 18 °C in a shaking incubator. Cells were harvested at 6,000 x g for 15 min at 4 °C and cell pellets were stored at −20 °C until purification. Cell pellets were resuspended in twice their volume of buffer A (BhcR: 100 mM KCl, 20 mM HEPES-KOH pH 7.5, 10 mM MgCl_2_, 4 mM β-mercaptoethanol, 5% glycerol and one tablet of SIGMAFAST™ protease inhibitor cocktail, EDTA-free per L; GlcR: 500 mM NaCl, 20 mM Tris pH 8.0, 15 mM imidazole, 1 mM β-mercaptoethanol, 5% glycerol and one tablet of SIGMAFAST™ protease inhibitor cocktail, EDTA-free per L). The cell suspension was treated with a Sonopuls GM200 sonicator (BANDELIN electronic GmbH & Co. KG, Berlin, Germany) at an amplitude of 50% to lyse the cells and subsequently centrifuged at 50,000 x g and 4 °C for 1 h. The filtered supernatant (0.45 µm filter; Sarstedt, Nümbrecht, Germany) was loaded onto Protino® Ni-NTA Agarose (Macherey-Nagel, Düren, Germany) in a gravity column, which had previously been equilibrated with 5 column volumes of buffer A. The column was washed with 20 column volumes of buffer A and 5 column volumes of 85% buffer A and 15% buffer B and the His-tagged protein was eluted with buffer B (buffer A with 500 mM imidazole). The eluate was desalted using PD-10 desalting columns (GE Healthcare, Chicago, USA) and buffer C (BhcR: 100 mM KCl, 20 mM HEPES-KOH pH 7.5, 10 mM MgCl_2_, 5% glycerol and 1 mM DTT; GlcR: 100 mM NaCl, 20 mM Tris pH 8.0, 1 mM DTT, 5% glycerol). This was followed by purification on a size exclusion column (Superdex™ 200 pg, HiLoad™ 16/600; GE Healthcare, Chicago, USA) connected to an ÄKTA Pure system (GE Healthcare, Chicago, USA) using buffer C. 2 mL concentrated protein solution was injected, and flow was kept constant at 1 mL min^-1^. Elution fractions containing pure protein were determined via SDS-PAGE analysis (39) on 12.5 % gels. Purified proteins in buffer C were subsequently used for downstream experiments.

For heterologous overproduction of MBP-GlcR, the corresponding plasmid encoding for the respective protein was first transformed into chemically competent *E. coli* BL21 AI cells. The cells were then grown on LB agar plates containing 34 µg mL^-1^ chloramphenicol at 37 °C overnight. A starter culture in selective LB medium was inoculated from a single colony on the next day and left to grow overnight at 37°C in a shaking incubator. The starter culture was used on the next day to inoculate an expression culture in selective TB medium with a starting OD_600_ of 0.05. The expression culture was grown at 37 °C in a shaking incubator to an OD_600_ of 1.0, induced with 0.5 mM IPTG and 0.025% L-arabinose and subsequently grown overnight at 20 °C in a shaking incubator. Cells were harvested at 4,000 x g for 20 min at 4 °C and cell pellets were stored at −70 °C until purification. Cell pellets were resuspended in twice their volume of buffer A (50 mM HEPES pH 7.5, 500 mM KCl) with 5 mM MgCl_2_ and DNase I (Roche, Basel, Switzerland). The cell suspension was treated with a Sonopuls GM200 sonicator (BANDELIN electronic GmbH & Co. KG, Berlin, Germany) at an amplitude of 50% in order to lyse the cells and subsequently centrifuged at 100,000 x g and 4 °C for 45 min. The filtered supernatant (0.45 µm filter; Sarstedt, Nümbrecht, Germany) was loaded onto a Ni-NTA column (HisTrap HP 1 mL, Cytiva, Marlborough, USA) using the fast protein liquid chromatography (FPLC) system (Äkta Start, Cytiva). The system had previously been equilibrated with buffer A + 25 mM imidazole. The column was washed with buffer A and 75 mM imidazole, and MBP-GlcR was eluted with buffer A + 500 mM imidazole. The eluate was desalted using a HiTrap desalting column (Sephadex G-25 resin, Cytiva) and protein elution buffer (25 mM Tris-HCl pH 7.4, 100 mM NaCl).

### Genetic modification of *P. denitrificans*

Transfer of replicative plasmids into *P. denitrificans* was performed via conjugation using *E. coli* ST18 as donor strain according to previously described methods (14). Selection of conjugants was performed at 30 °C on LB plates containing 0.5 µg ml^-1^ tetracycline. Successful transfer of plasmids into *P. denitrificans* was verified by colony PCR.

Transfer of gene deletion plasmids into *P. denitrificans* was performed in the same way. Selection of conjugants was performed at 30 °C on LB agar plates containing 25 µg mL^-1^ kanamycin. The respective gene deletion was verified by colony PCR and DNA sequencing (Eurofins Genomics, Ebersberg, Germany), and the deletion strain was propagated in selective LB medium. In each case, the gene to be deleted was replaced by a kanamycin resistance cassette either in the same direction or the opposite direction to exclude polar effects.

### High-throughput growth and fluorescence assays with *P. denitrificans* strains

Cultures of *P. denitrificans* DSM 413 WT and its derivatives were pre-grown at 30 °C in LB medium containing 25 µg mL^-1^ kanamycin or 0.5 µg mL^-1^ tetracycline, when necessary. Cells were harvested, washed once with minimal medium containing no carbon source and used to inoculate growth cultures of 180 µL minimal medium containing an appropriate carbon source as well as 25 µg mL^-1^ kanamycin or 0.5 µg mL^-1^ tetracycline, when necessary. Growth and fluorescence in 96-well plates (Thermo Fisher Scientific, Waltham, USA) were monitored at 30 °C at 600 nm in a Tecan Infinite M200Pro plate reader (Tecan, Männedorf, Switzerland). Fluorescence of mCherry was measured at an emission wavelength of 610 nm after excitation at 575 nm. The resulting data was evaluated using GraphPad Prism 8.1.1. To determine whether differences in growth rate or substrate uptake rate are significant, unpaired t tests with Welch’s correction were used.

### Whole-cell shotgun proteomics

To acquire the proteome of *P. denitrificans* WT and Δ*cceR* (OD***_600_ ∼***0.4) in minimal medium supplemented with 60 mM glyoxylate, four replicate cultures were grown for each strain. Main cultures were inoculated from precultures grown in the same medium in a 1:1,000 dilution. Cultures were harvested by centrifugation at 4,000 **×** g and 4 **°**C for 15 min. Supernatant was discarded and pellets were washed in 40 mL phosphate buffered saline (PBS; 137 mM NaCl, 2.7 mM KCl, 10 mM Na***_2_***HPO***_4_***, 1.8 mM KH***_2_***PO***_4_***, pH 7.4). After washing, cell pellets were resuspended in 1 mL PBS, transferred into Eppendorf tubes, and repeatedly centrifuged. Cell pellets in Eppendorf tubes were snap-frozen in liquid nitrogen and stored at **-**80 **°**C until they were used for the preparation of samples for LC-MS analysis and label-free quantification.

For protein extraction bacterial cell pellets were resuspended in 4% sodium dodecyl sulfate (SDS) and lysed by heating (95 °C, 15 min) and sonication (Hielscher Ultrasonics GmbH, Teltow, Germany). Reduction was performed for 15 min at 90 °C in the presence of 5 mM tris(2-carboxyethyl)phosphine followed by alkylation using 10 mM iodoacetamide at 25 °C for 30 min. The protein concentration in each sample was determined using the BCA protein assay kit (Thermo Fisher Scientific, Waltham, USA) following the manufacturer’s instructions. Protein cleanup and tryptic digest were performed using the SP3 protocol as described previously (40) with minor modifications regarding protein digestion temperature and solid phase extraction of peptides. SP3 beads were obtained from GE Healthcare (Chicago, USA). 1 µg trypsin (Promega, Fitchburg, USA) was used to digest 50 μg of total solubilized protein from each sample. Tryptic digest was performed overnight at 30 °C. Subsequently, all protein digests were desalted using C18 microspin columns (Harvard Apparatus, Holliston, USA) according to the manufacturer’s instructions.

LC-MS/MS analysis of protein digests was performed on a Q-Exactive Plus mass spectrometer connected to an electrospray ion source (Thermo Fisher Scientific, Waltham, USA). Peptide separation was carried out using an Ultimate 3000 nanoLC-system (Thermo Fisher Scientific, Waltham, USA), equipped with an in-house packed C18 resin column (Magic C18 AQ 2.4 µm; Dr. Maisch, Ammerbuch-Entringen, Germany). The peptides were first loaded onto a C18 precolumn (preconcentration set-up) and then eluted in backflush mode with a gradient from 94% solvent A (0.15% formic acid) and 6% solvent B (99.85% acetonitrile, 0.15% formic acid) to 25% solvent B over 87 min, continued with 25% to 35% of solvent B for an additional 33 min. The flow rate was set to 300 nL/min. The data acquisition mode for the initial LFQ study was set to obtain one high-resolution MS scan at a resolution of 60,000 (*m/z* 200) with scanning range from 375 to 1500 *m/z* followed by MS/MS scans of the 10 most intense ions. To increase the efficiency of MS/MS shots, the charged state screening modus was adjusted to exclude unassigned and singly charged ions. The dynamic exclusion duration was set to 30 sec. The ion accumulation time was set to 50 ms (both MS and MS/MS). The automatic gain control (AGC) was set to 3 × 10^6^ for MS survey scans and 1 × 10^5^ for MS/MS scans. Label-free quantification was performed using Progenesis QI (version 2.0). MS raw files were imported into Progenesis and the output data (MS/MS spectra) were exported in mgf format. MS/MS spectra were then searched using MASCOT (version 2.5) against a database of the predicted proteome from *P. denitrificans* downloaded from the UniProt database (www.uniprot.org; download date 01/26/2017), containing 386 common contaminant/background proteins that were manually added. The following search parameters were used: full tryptic specificity required (cleavage after lysine or arginine residues); two missed cleavages allowed; carbamidomethylation (C) set as a fixed modification; and oxidation (M) set as a variable modification. The mass tolerance was set to 10 ppm for precursor ions and 0.02 Da for fragment ions for high energy-collision dissociation (HCD). Results from the database search were imported back to Progenesis, mapping peptide identifications to MS1 features. The peak heights of all MS1 features annotated with the same peptide sequence were summed, and protein abundance was calculated per LC–MS run. Next, the data obtained from Progenesis were evaluated using the SafeQuant R-package version 2.2.2 (41).

### Electrophoretic mobility shift assays

Fluorescently labeled DNA fragments for electrophoretic mobility shift assays (EMSA) were generated by PCR from genomic DNA of *P. denitrificans* DSM 413. For the *Pbhc* regulatory region, primers Pbhc_fw and Pbhc_rev-dye were used to generate a 238-bp fragment containing the putative *Pbhc* promoter. The primers bhcA_fw and bhcA_rev-dye were used to generate a 255-bp fragment containing a part of the *bhcA* gene as negative control. For the *Pglc* regulatory region, primers Pglc_fw and Pglc_rev-dye were used to generate a 156-bp fragment containing the putative *Pglc* promoter.

The primers glcD_fw and glcD_rev-dye were used to generate a 156-bp fragment containing a part of the *glcD* gene as negative control. All respective reverse primers were 5’-labelled with the Dyomics 781 fluorescent dye (Microsynth AG, Balgach, Switzerland). Binding reactions were performed in buffer A (20 mM potassium phosphate pH 7.0, 1 mM DTT, 5 mM MgCl_2_, 50 mM KCl, 15 µg mL^-1^ bovine serum albumin, 50 µg mL^-1^ herring sperm DNA, 5% v/v glycerol, 0.1% Tween20) in a total volume of 20 µL. The respective DNA fragments (0.025 pM) were incubated with various amounts of the purified protein BhcR (0x/400x/2,000x/4,000x/10,000x/20,000x/30,000x/40,000x molar excess) or GlcR (0x/ 20x/100x/200x/500x/1,000x/1,500x/2,000x molar excess) and protein:DNA complexes were incubated with various concentrations of effector molecules as indicated in the respective figure legends. After incubation of the reaction mixtures at 37 °C for 20 min, the samples were loaded onto a native 5% polyacrylamide gel and electrophoretically separated at 110 V for 60 min. BhcR/GlcR:DNA-interactions were detected using an Odyssey FC Imaging System (LI-COR Biosciences, Lincoln, USA).

### Fluorescence polarization assays

Fluorescently labeled DNA fragments for fluorescence polarization assays were generated as follows: 1) For the [6FAM]-*P_glc_* fragment, DNA was amplified from pTE714_4400/4399_ig using primers [6FAM]-Pglc_fw & Pglc_rev. 2) For [6FAM]-*tetO*, 10 µM [6FAM]-tetO_fw and 10 µM tetO_rev primers were mixed in 1x annealing buffer (15 mM phosphate buffer pH 7.3, 0.5 mM EDTA, 7 mM MgCl_2_, 0.01% Triton X-100). Primers were annealed by incubation at 95 °C for 2 min and subsequent cool-down to room temperature in the heating block. All fluorescence polarization experiments were prepared in 1x binding buffer (50 mM HEPES pH 7.5, 100 mM NaCl, 0.01% Triton X-100) using 10 nM DNA at 20 µL scale. All reagents were prepared in 1x binding buffer. For the GlcR binding curve, the MGlcR dilution series and 20 nM DNA dilutions ([6FAM]-*P_glc_* and [6FAM]-*tetO*) were mixed at equal volumes of 10 µL. MGlcR was assayed in a 1:2 dilution series from 6 µM to 5.86 nM. For the effector binding assay, 10 µL MGlcR at 1.5 µM, 5 µL [6FAM]-*P_glc_*DNA at 40 nM and 5 µL respective effector dilution series were mixed. Effectors were assayed at 100, 10, 1 and 0.1 mM. Reactions were transferred into black, non-binding 384-well plates (Greiner BioOne, Kremsmünster, Austria), briefly centrifuged in a benchtop centrifuge, and incubated for 10 min at room temperature. Fluorescence polarization was measured in a Tecan Spark (Tecan, Männedorf, Switzerland) with 485 nm excitation and 535 nm emission wavelength, using optimal gain and optimal Z-position as determined by the plate reader. Blanks were prepared with 1x binding buffer.

### Substrate uptake experiments

Quantitative determination of glycolate in spent medium was performed using a LC-MS/MS. The chromatographic separation was performed on an Agilent Infinity II 1290 HPLC system using a Kinetex EVO C18 column (150 × 1.7 mm, 3 μm particle size, 100 Å pore size, Phenomenex) connected to a guard column of similar specificity (20 × 2.1 mm, 5 μm particle size, Phenomenex) at a constant flow rate of 0.1 ml/min with mobile phase A being 0.1% formic acid in water and phase B being 0.1% formic acid in methanol (Honeywell, Morristown, New Jersey, USA) at 25 °C.

The injection volume was 1 µl. The mobile phase profile consisted of the following steps and linear gradients: 0 – 4 min constant at 0 % B; 4 – 6 min from 0 to 100 % B; 6 – 7 min constant at 100 % B; 7 –7.1 min from 100 to 0 % B; 7.1 to 12 min constant at 0 % B. An Agilent 6495 ion funnel mass spectrometer was used in negative mode with an electrospray ionization source and the following conditions: ESI spray voltage 2000 V, nozzle voltage 500 V, sheath gas 400 °C at 11 l/min, nebulizer pressure 50 psig and drying gas 80 °C at 16 l/min. The target compound was identified based on its mass transitions and retention time compared to standards. Chromatograms were integrated using MassHunter software (Agilent, Santa Clara, California, USA). Absolute concentrations were calculated based on an external calibration curve prepared in fresh medium. Mass transitions, collision energies, cell accelerator voltages and dwell times were optimized using chemically pure standards. Parameter settings for glycolate were as follows: quantifier 75→75; collision energy 0; qualifier 75→47; collision energy 6; dwell 20; cell accelerator voltage 5.

Glucose concentrations in spent medium were quantified using the Glucose-Glo™ Assay kit (Promega, Walldorf, Germany). Luminescence measurements of diluted medium samples were performed in white 384-well plates (Greiner BioOne, Kremsmünster, Austria) in a Tecan Infinite M200Pro plate reader (Tecan, Männedorf, Switzerland) according to the instructions of the kit.

### Phylogenetic analyses

Sequences of BhcR homologs and other transcriptional regulators of the IclR family were downloaded from the NCBI Protein database (https://www.ncbi.nlm.nih.gov/protein/) and aligned using MUSCLE (42). A maximum likelihood phylogenetic tree of the aligned sequences was calculated with MEGA X (43) using the Le-Gascuel model (44) with 100 bootstraps. The resulting tree was visualized using iTOL (45). The phylogenetic tree for GlcR homologs and other transcriptional regulators of the FadR subfamily was generated in the same way.

The alignment of IclR, AllR, and BhcR amino acid sequences was generated using MUSCLE and colored with Jalview (46).

### Visualization and statistical analysis

Data were evaluated and visualized using GraphPad Prism 8.1.1., and results were compared using an unpaired t-test with Welch’s correction in GraphPad Prism 8.1.1.

## Results

### Glyoxylate assimilation via the BHAC is regulated by BhcR

We first focused on understanding the regulation of the BHAC, which mediates the second step of glycolate metabolism in *P. denitrificans*. To investigate the role of the transcription factor BhcR in regulating this pathway, we characterized the protein bioinformatically and experimentally. Amino acid sequence analysis showed that BhcR contains an IclR-type helix-turn-helix domain and an IclR-type effector-binding domain (see Uniprot: https://www.uniprot.org/uniprotkb/A1B8Z4), indicating that the protein belongs to the IclR-type family of transcriptional regulators. In a phylogenetic tree of 1083 sequences from 29 subfamilies within the IclR-type family, BhcR formed a close sister group to a clade of IclR and AllR homologs (**Supplementary Figure 1**).

IclR, the namesake representative of the family, regulates expression of the glyoxylate shunt operon (*aceBAK*) in *E. coli* and other bacteria. The protein forms a tetramer that acts as transcriptional repressor. IclR is allosterically regulated by glyoxylate and pyruvate, which control the oligomerization state of IclR. Pyruvate stabilizes tetramer formation, while glyoxylate favors dimer formation and releases IclR from the DNA (47). AllR acts as transcriptional repressor of the allantoin and glyoxylate utilization operons in *E. coli*. It binds to the *gcl* promoter and the *allS*-*allA* intergenic region. Similarly to IclR, DNA-binding of AllR is decreased by increasing concentrations of glyoxylate (48).

In IclR, glyoxylate and pyruvate occupy the same binding site. With the exception of one residue, this ligand-binding site is conserved in AllR (**Table 1, Supplementary Figure 2**), while in BhcR the putative binding site shows some marked differences. Amino acids that bind to the oxygen atoms of glyoxylate or pyruvate are conserved between IclR and BhcR (except for the presence of isoleucine in place of alanine at position 161). In contrast, a hydrophobic patch of residues that interacts with the methyl group of pyruvate in IclR is apparently lacking in BhcR (**Table 1**).

**Table 1:** Ligand-binding residues of IclR family transcriptional regulators. Ligand-binding residues of *E. coli* IclR that were previously described (47) are compared to their counterparts in *E. coli* AllR and *P. denitrificans* BhcR. Numbering is based on the sequence of *E. coli* IclR.

|  | IclR | AllR | BhcR |
| --- | --- | --- | --- |
| **residues in hydrophobic patch that interact with methyl group of pyruvate** |  |  |  |
| **143** | Leu | Met | Thr |
| **146** | Met | Met | Ala |
| **154** | Leu | Leu | Ser |
| **220** | Leu | Leu | Met |
| **residues that bind to oxygen atoms of glyoxylate or pyruvate** |  |  |  |
| **160** | Gly | Gly | Gly |
| **161** | Ala | Ala | Ile |
| **212** | Asp | Asp | Asp |
| **239** | Ser | Ser | Ser |
| **241** | Ser | Ser | Ser |

To study BhcR in more detail, we purified the regulator from *P. denitrificans* and conducted additional DNA binding experiments with the putative promoter region of the *bhc* gene cluster (P*_bhc_*) (**Figure 1a**). This region contains a palindromic sequence close to the potential −35 region of the *bhcABCD* gene cluster, which could serve as potential binding site for BhcR (**Figure 1b**). In electrophoretic mobility shift assays (EMSAs), the interaction of BhcR with P*_bhc_* was negatively affected by increasing concentrations of glyoxylate, as previously described (14). In contrast, DNA-binding interaction was positively affected by the presence of pyruvate or oxalate (**Figure 1a**), suggesting that these two molecules stabilize the tetrameric DNA-binding form of BhcR, analogous to the reported interaction of pyruvate with IclR (47). Interestingly, *P. denitrificans* was not capable of growth on oxalate as sole source of carbon and energy (**Supplementary Figure 3**), indicating that the observed *in vitro* interaction of BhcR with this compound might not be relevant *in vivo*.

**Figure 1:**
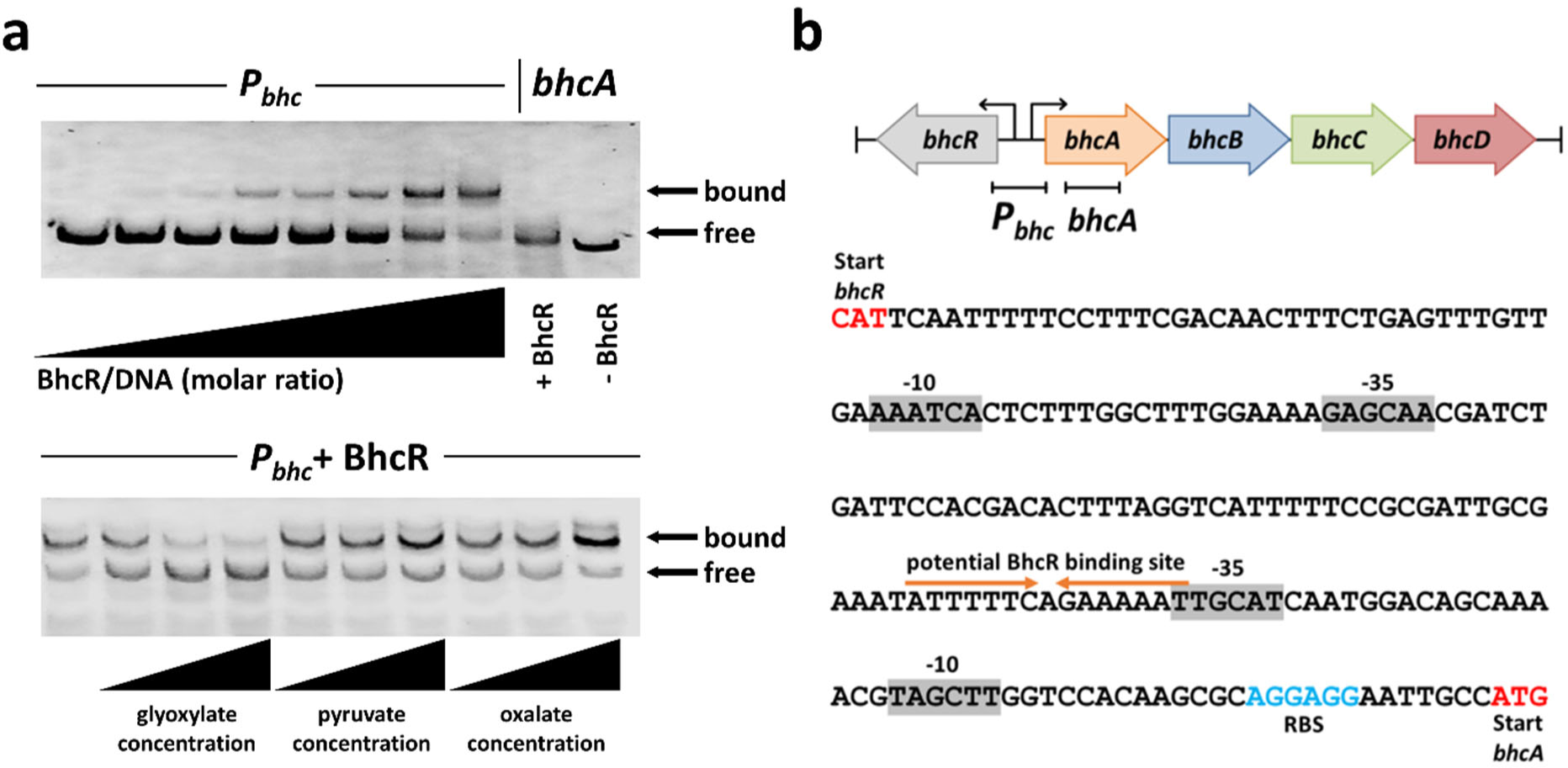
DNA-binding properties of BhcR. **a**, Top, a fluorescently labelled 238 bp DNA fragment carrying the putative promoter region of the *bhc* gene cluster (*P_bhc_*) was incubated with increasing amounts of purified BhcR protein (0x/400x/2,000x/4,000x/10,000 x/20,000 x/30,000x/40,000x molar excess) and subsequently separated by electrophoresis to visualize DNA bound to BhcR and free DNA; a 255 bp DNA fragment derived from the coding region of *bhcA* was used as a negative control. BhcR specifically forms a complex with the *P_bhc_* DNA fragment. Bottom, the *P_bhc_*–BhcR complex (40,000x molar excess BhcR) was incubated with increasing concentrations (0.1 mM; 0.5 mM; 5 mM) of glyoxylate, pyruvate, or oxalate, and subsequently separated by electrophoresis to assess the effect of these metabolites on complex formation. Increasing concentrations of glyoxylate decrease the binding of BhcR to the *P_bhc_*DNA fragment, while the opposite effect is observed for increasing concentrations of pyruvate or oxalate. **b**, DNA binding of BhcR in the P*_bhc_*promoter region. Potential −35 and −10 regions upstream of the *bhcR* and *bhcA* genes were identified using BPROM (49). A potential palindromic binding site for BhcR was identified upstream of the −35 region of *bhcA*.

Next, we generated a *P. denitrificans* Δ*bhcR* deletion strain and tested its growth on different carbon sources. In this strain, *bhcR* was replaced by a kanamycin resistance cassette in the same transcriptional direction. We also created a control strain, in which we inserted the kanamycin cassette in the opposite transcriptional direction to exclude polar effects. Both deletion strains were unable to grow on glycolate or glyoxylate (**Figure 2a+b**), while growth on acetate, succinate, pyruvate or glucose was not affected (**Figure 2c**). A similar phenotype was recently observed for a *bhcABCD* deletion strain (14), which suggests that BhcR acts as an activator that is required for transcription of the *bhc* gene cluster. We sought to further investigate this hypothesis by generating P*_bhc_* promoter-based reporter strains with mCherry as reporter. We tested mCherry production in the Δ*bhcR*, Δ*bhcABCD*, and wild-type strain (**Figure 2d**). When grown on succinate, only low fluorescence levels were observed in all three strains, indicating a basal expression of the *bhc* gene cluster. Supplementation of succinate medium with increasing concentrations of glyoxylate caused a gradual increase in fluorescence in the WT and Δ*bhcABCD* backgrounds, suggesting an increase in P*_bhc_* promoter activity. Notably, in these experiments, promoter activity was positively correlated with the intracellular concentration of glyoxylate. The Δ*bhcABCD* strain that cannot further convert glyoxylate (resulting in higher intracellular glyoxylate levels) exhibited significantly higher expression from the P*_bhc_* promoter compared to the WT strain, in which glyoxylate is continuously converted via the BHAC. In contrast to these two strains, expression from the P*_bhc_* promoter remained basal in the Δ*bhcR* background even in the presence of glyoxylate, supporting the role of BhcR as activator of the *bhc* gene cluster *in vivo*.

**Figure 2:**
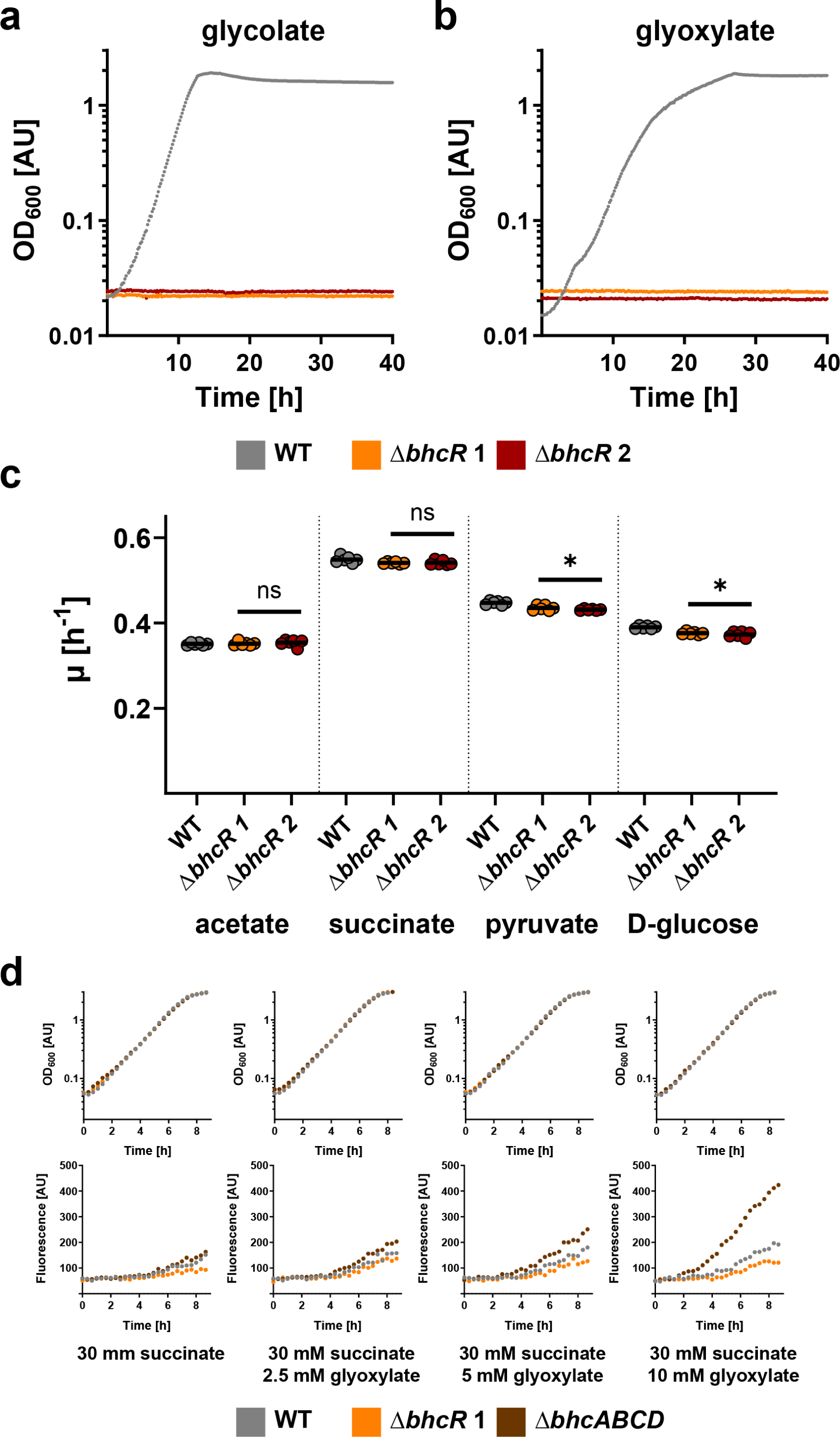
Characterization of *P. denitrificans* Δ*bhcR*. **a**, **b**, Growth curves of wild-type *P. denitrificans* DSM 413 (grey) and *bhcR* deletion strains (orange + red) grown in the presence of 60 mM glycolate (**a**) or 60 mM glyoxylate (**b**). Deletion of *bhcR* is sufficient to abolish growth in the presence of these carbon sources. These experiments were repeated three times independently with similar results. **c**, Growth rates (*μ*) of wild-type *P. denitrificans* DSM 413 (grey) and *bhcR* deletion strains (orange + red) grown in the presence of 60 mM acetate, 30 mM succinate, 40 mM pyruvate, or 20 mM glucose. The growth rates of the *bhcR* deletion strains were either not significantly changed or only slightly decreased on these substrates when compared to the wild-type. The results of *n* = 6 independent experiments are shown, and the black line represents the mean. **d**, Growth and fluorescence of promoter reporter strains Δ*bhcR* (orange), Δ*bhcABCD* (brown), and WT (grey) with pTE714-*P_bhc_* on different carbon sources. These experiments were repeated three times independently with similar results. Growth and fluorescence of negative control strains are shown in **Supplementary Figure 4**.

How can the *in vivo* function of BhcR as activator of P*_bhc_* be reconciled with the *in vitro* data that showed decreased DNA binding in the presence of glyoxylate? The most likely possibility is that BhcR also represses its own expression in the absence of glyoxylate, but activates the expression of the *bhc* gene cluster in the presence of glyoxylate. This dual function would explain the decreased *in vitro* DNA binding of BhcR in the presence of glyoxylate. Notably, such a dual role as activator and repressor was previously described for other IclR family regulators (50, 51) and for the transcriptional activator/repressor RamB, a member of the ScfR family in *P. denitrificans* (32).

### Pden_4400 encodes for GlcR, a novel repressor of the glycolate oxidase gene cluster

Next, we studied the regulation of glycolate oxidation in *P. denitrificans*. We hypothesized that glycolate is converted into glyoxylate by the three-subunit enzyme glycolate oxidase (GlcDEF), encoded by the genes Pden_4397-99, and verified the role of this gene cluster by generating a Pden_4397-99 deletion strain, which was unable to grow on glycolate as sole carbon source (**Supplementary Figure 5**). The gene Pden_4400, adjacent to this gene cluster, is annotated as a transcriptional regulator of the GntR family. This resembles the situation in *E. coli*, where the GntR-family regulator GlcC serves as transcriptional activator of *glcDEF* (5, 52). GlcC is part of the FadR subfamily of the GntR transcription factor family (53). In a phylogenetic tree containing sequences of GlcC homologs, Pden_4400 homologs, and sequences from other clades within the FadR subfamily (283 sequences in total; **Supplementary Figure 6**), Pden_4400 and its close homologs form a well-defined clade that clusters together with the GlcC clade, as well as the PdhR (regulator of pyruvate dehydrogenase (54)), and LldR (regulator of lactate dehydrogenase (55, 56)) clades. This suggests that Pden_4400 might fulfill a similar role as GlcC, but is not simply an alphaproteobacterial homolog of this transcriptional activator. We therefore designate Pden_4400 as *glcR*.

Homologs of *glcR* can be found adjacent to *glcDEF* in many *Paracoccus* strains, but also in other *Rhodobacterales* (e.g., *Methylarcula*, *Puniceibacterium*, *Rhodobacter*) as well as in some *Rhizobiales* (e.g., *Afipia*, *Chenggangzhangella*) (**Supplementary Table 1**), suggesting that control of glycolate oxidase production via GlcR is conserved across different alphaproteobacterial clades.

To study GlcR in more detail, we purified the transcriptional regulator and investigated its DNA-binding capabilities. In EMSAs, we could demonstrate specific binding of the protein to a DNA fragment containing the putative promoter region of the *glc* gene cluster (*P_glc_*). DNA binding was decreased in the presence of glycolate, while glyoxylate did not alter DNA binding of GlcR (**Figure 3a+b**). We subsequently purified GlcR fused to an N-terminal maltose-binding protein (MGlcR) to increase its solubility for fluorescence polarization experiments. These experiments confirmed previous results with the non-tagged protein, and allowed us to determine a K_D_ for MGlcR of 225 ± 5 nM at 10 nM DNA. Titration of the *P_glc_*-MGlcR complex with increasing concentrations of glycolate demonstrated a notable decrease in binding, while the same effect was not observed for glyoxylate (**Figure 3c+d**).

**Figure 3:**
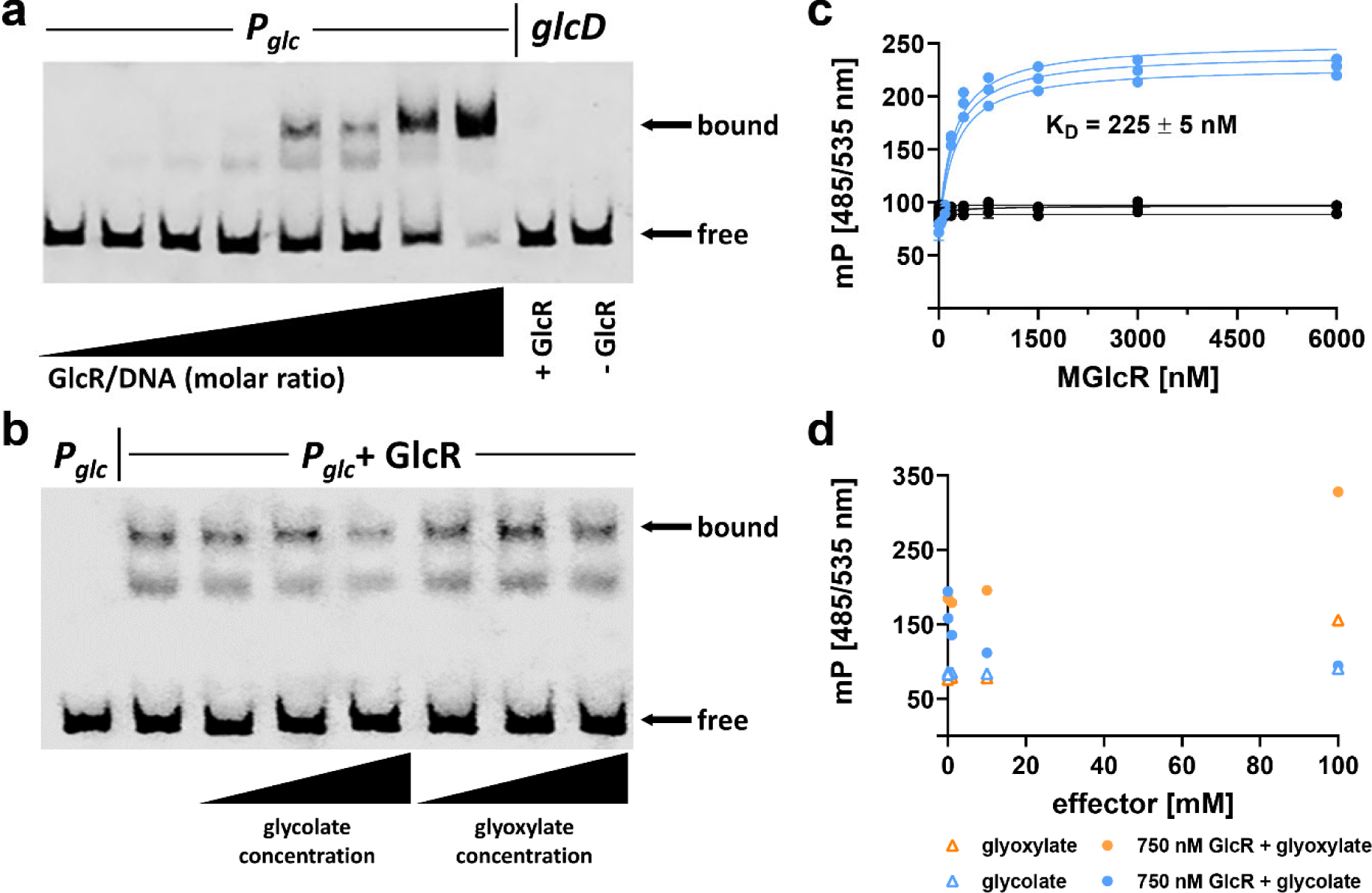
DNA-binding properties of GlcR. **a**, a fluorescently labelled 156 bp DNA fragment carrying the putative promoter region of the *glc* gene cluster (*P_glc_*) was incubated with increasing amounts of purified GlcR protein (0x/20x/100x/200x/500x/1,000x/1,500x/2,000x molar excess) and subsequently separated by electrophoresis to visualize DNA bound to GlcR and free DNA; a 156 bp DNA fragment derived from the coding region of *glcD* was used as a negative control. GlcR specifically forms a complex with the *P_glc_*DNA fragment. **b**, the *P_glc_*–GlcR complex (1,000x molar excess of GlcR) was incubated with increasing concentrations of glycolate or glyoxylate (0.5 mM, 1 mM, 2 mM) and subsequently separated by electrophoresis to assess the effect of these metabolites on DNA:GlcR complex formation. Increasing concentrations of glycolate decrease the binding of GlcR to *P_glc_*, while increasing concentrations of glyoxylate did not result in altered DNA binding of GlcR. **c**, fluorescence polarization experiments with increasing concentrations of MGlcR and the *P_glc_* region (blue) or the *tetO* sequence as negative control (black). Three independent experiments were conducted for each combination, and a K_D_ of 225 ± 5 nM was determined for MGlcR with 10 nM *P_glc_*. **d**, fluorescence polarization experiments with increasing concentrations of an effector (glycolate or glyoxylate; 0, 0.1, 1, 10, 100 mM) and 750 nM MGlcR and 10 nM *P_glc_*. These results confirm that glycolate causes decreased binding of GlcR to *P_glc_*.

Subsequently, we generated two *P. denitrificans* deletion strains of *glcR* and tested their growth on different carbon sources. As for *bhcR*, the *glcR* gene was replaced with a kanamycin resistance cassette in either the same or the opposite direction of transcription to exclude any polar effects. Interestingly, the growth rate of the deletion strains on glycolate was not significantly different from the WT (**Figure 4a+c**). However, the growth rates on glyoxylate, but also on succinate and acetate, were slightly decreased compared to the WT (**Figure 4b+c**). Taken together, these data strongly suggest that GlcR does not act as activator, but as repressor. In the *glcR* deletion strain, GlcDEF is constitutively produced, which explains the WT-like behavior of the deletion strain on glycolate, and the slightly decreased growth rate of the deletion strain on glyoxylate, succinate, and acetate due to increased protein production burden.

**Figure 4:**
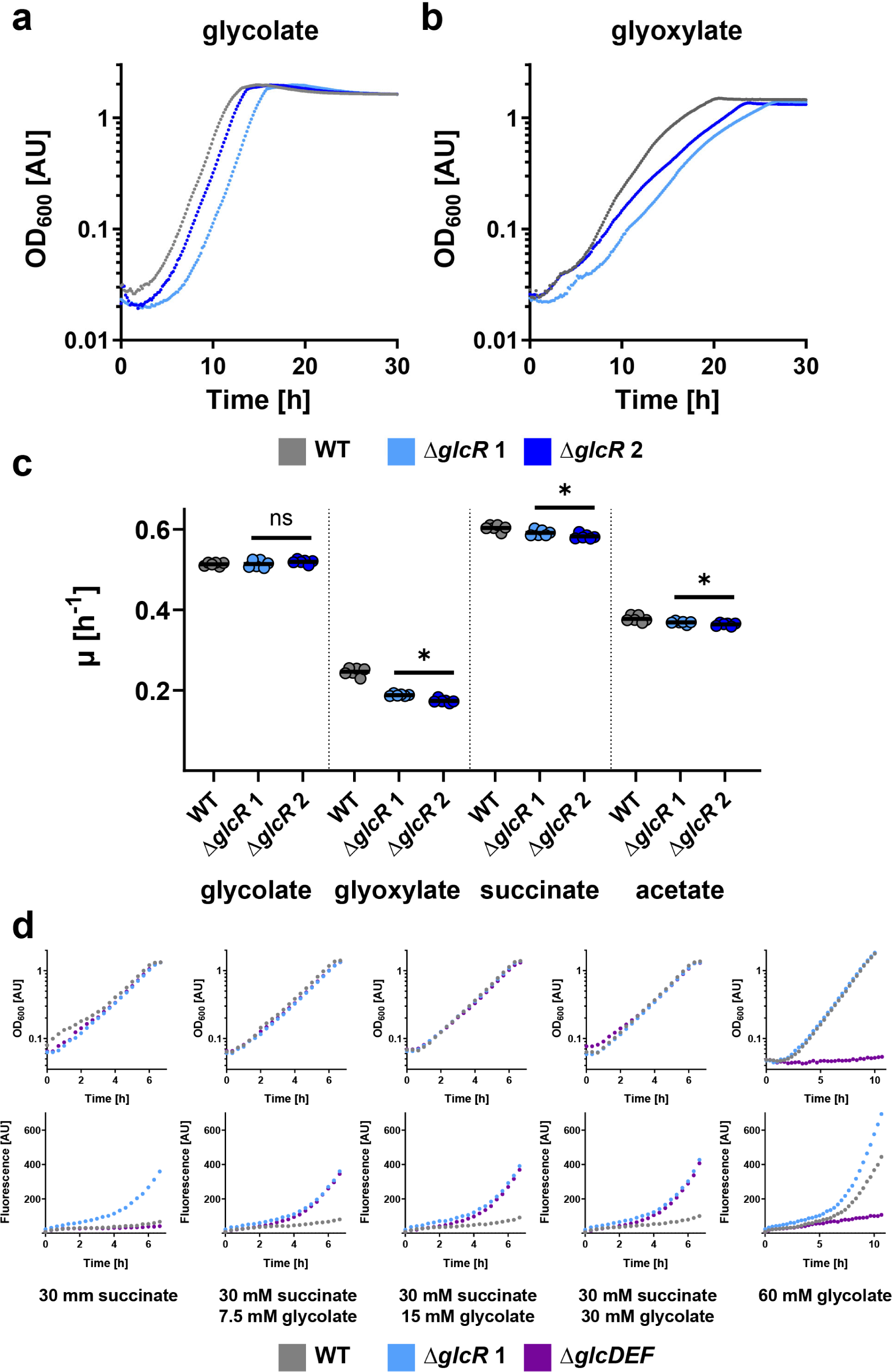
Characterization of *P. denitrificans* Δ*glcR*. **a**, **b**, Growth curves of wild-type *P. denitrificans* DSM 413 (grey) and *glcR* deletion strains (light + dark blue) grown in the presence of 60 mM glycolate (**a**) or 60 mM glyoxylate (**b**). These experiments were repeated three times independently with similar results. **c**, Growth rates (*μ*) of wild-type *P. denitrificans* DSM 413 (grey) and *glcR* deletion strains (light + dark blue) grown in the presence of 60 mM glycolate, 60 mM glyoxylate, 30 mM succinate, or 60 mM acetate. When compared to the wild-type, the growth rates of the *glcR* deletion strains were slightly decreased in the presence of glyoxylate, succinate, and acetate. The results of *n* = 6 independent experiments are shown, and the black line represents the mean. **d**, Growth and fluorescence of promoter reporter strains Δ*glcR* (light blue), Δ*glcDEF* (purple), and WT (grey) with pTE714-*P_glc_* on different carbon sources. These experiments were repeated three times independently with similar results. Growth and fluorescence of negative control strains are shown in **Supplementary Figure 4**.

We independently confirmed the role of GlcR in *P. denitrificans* using P*_glc_* promoter-based reporter strains. We tested under which conditions mCherry was produced from a P*_glc_*-fusion in the Δ*glcR*, Δ*glcDEF*, and WT background (**Figure 4d**). When growing on succinate, fluorescence only increased in the Δ*glcR* background, but not in the other two strains. This increase is consistent with the finding that GlcR acts as repressor *in vitro*. When growing on succinate and different concentrations of glycolate, fluorescence also increased in the Δ*glcDEF* background, but only slightly in the WT. This can be explained by the fact that in the WT intracellular glycolate levels stay relatively low, as glycolate is further metabolized. In contrast, glycolate accumulates in the Δ*glcDEF* strain, which is incapable of converting glycolate further to glyoxylate due to the lack of glycolate oxidase, resulting in increased expression from *P_glc_*. Finally, with glycolate as sole carbon source, fluorescence also increased in the WT background (while the Δ*glcDEF* strain was unable to grow under these conditions).

### Growth of *P. denitrificans* on two carbon substrates does not result in diauxie

Having characterized the regulatory circuits of glycolate oxidase and the BHAC at the molecular level, we aimed at studying glycolate and glyoxylate metabolism under more complex growth conditions at the cellular level. To that end, we grew *P. denitrificans* on glycolate (or glyoxylate) together with either glucose, a glycolytic carbon substrate, or pyruvate, a gluconeogenic carbon substrate, to determine the effect of substrate co-feeding on growth.

We first grew *P. denitrificans* either on a single carbon substrate or on two carbon substrates, mixed in three different ratios. Growth on glycolate (µ = 0.51 h^-1^) was faster than growth on glyoxylate (0.28 h^-1^), while the growth rates on pyruvate (µ = 0.45 h^-1^) and glucose (µ = 0.38 h^-1^) were between these two values. When growing on a mix of glycolate and glucose (**Figure 5a**), the growth rate of *P. denitrificans* was not different from the growth rate on glycolate alone, while the growth rate of the bacterium was very similar to the growth rate on glucose alone when growing on a mix of glyoxylate and glucose (**Figure 5b**). The same pattern was also observed when glucose was replaced with pyruvate (**Figure 5c+d**). Notably, we did not observe any diauxic growth behavior (i.e., a first growth phase, an intermediate lag phase, and a second growth phase) on any of the tested carbon substrate mixtures.

**Figure 5:**
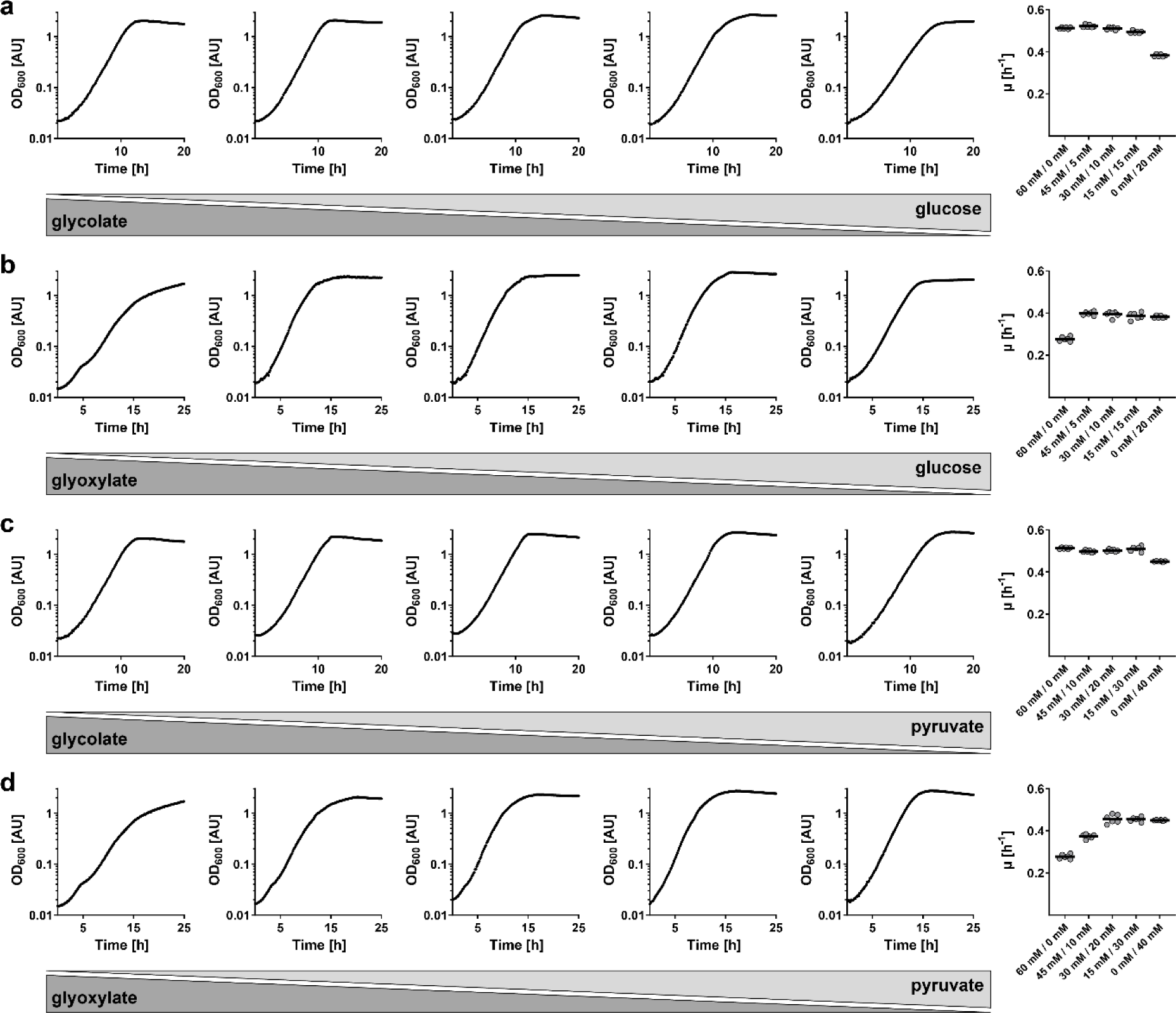
Growth of *P. denitrificans* on two carbon substrates. **a,** Growth on different concentrations of glycolate and glucose (from left to right: 60 mM/0 mM, 45 mM/5 mM, 30 mM/10 mM, 15 mM/15 mM, 0 mM/20 mM). **b,** Growth on different concentrations of glyoxylate and glucose (from left to right: 60 mM/0 mM, 45 mM/5 mM, 30 mM/10 mM, 15 mM/15 mM, 0 mM/20 mM). **c,** Growth on different concentrations of glycolate and pyruvate (from left to right: 60 mM/0 mM, 45 mM/10 mM, 30 mM/20 mM, 15 mM/30 mM, 0 mM/40 mM). **d,** Growth on different concentrations of glyoxylate and pyruvate (from left to right: 60 mM/0 mM, 45 mM/10 mM, 30 mM/20 mM, 15 mM/30 mM, 0 mM/40 mM). On the right of each panel, average growth rates from *n* = 6 independent growth experiments are shown.

Collectively, these data suggested that *P. denitrificans* does not assimilate the two carbon substrates sequentially, but rather in a co-utilizing manner. We therefore set out to investigate the regulation of central carbon metabolism and the uptake hierarchy of carbon substrates in *P. denitrificans* in more detail, with a special focus on glycolate and glyoxylate.

### CceR regulates glycolysis and gluconeogenesis in *P. denitrificans*

To this end, we investigated the role of the transcription factor CceR (<u>c</u>entral <u>c</u>arbon and <u>e</u>nergy metabolism regulator) in glycolate and glyoxylate metabolism of *P. denitrificans*. This protein was previously described as key regulator of carbon and energy metabolism in the Alphaproteobacterium *Rhodobacter sphaeroides*. CceR was also identified in *P. denitrificans*, where it was predicted to share largely the same regulon as in *R. sphaeroides* (57). Specifically, we aimed to determine whether CceR controls glycolate/glyoxylate assimilation pathways, uptake of these substrates into the cell, or both. We therefore generated two *P. denitrificans* Δ*cceR* strains, in which the gene was replaced with a kanamycin resistance cassette in either the same or the opposite direction of transcription.

Subsequently, we determined the growth rates of the Δ*cceR* and WT strains on 21 different carbon sources, including glycolate and glyoxylate (**Figure 6**). Notably, Δ*cceR* strains had reduced growth rates on all gluconeogenic carbon sources, but not on the five glycolytic carbon sources. This partially contrasts the situation in *R. sphaeroides*, where the growth rates of Δ*cceR* were not significantly decreased on the gluconeogenic carbon sources acetate, tartrate, aspartate, and isoleucine (57), indicating few, but distinct differences in the regulation of central carbon metabolism between both bacteria.

**Figure 6:**
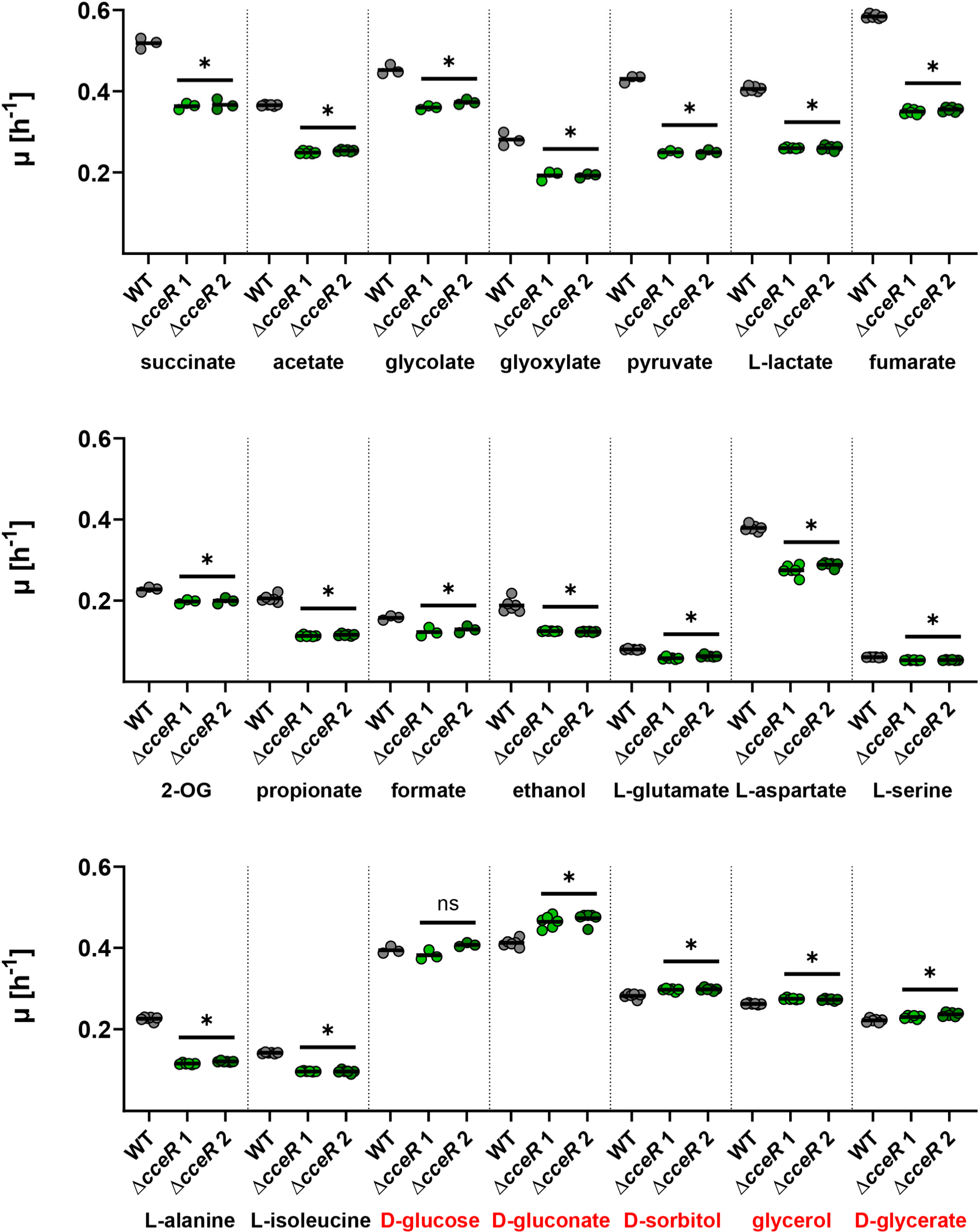
Characterization of *P. denitrificans* Δ*cceR*. Growth rates (*μ*) of wild-type *P. denitrificans* DSM 413 (grey) and *cceR* deletion strains (light + dark green) grown in the presence of various carbon sources (final carbon concentration 120 mM). When compared to the wild-type, the growth rates of the Δ*cceR* strains were significantly decreased in the presence of all substrates, except for the glycolytic carbon sources D-glucose, D-gluconate, D-sorbitol, glycerol, and D-glycerate (highlighted in red in the last row). The results of *n* ≥ 3 independent experiments are shown, and the black line represents the mean. 2-OG: 2-oxoglutarate.

We then analyzed the proteome of *P. denitrificans* WT and Δ*cceR* during growth on glyoxylate to identify the CceR regulon and its potential effects on C2 metabolism (**Figure 7**). Notably, several key enzymes of gluconeogenesis were downregulated in the Δ*cceR* strain, including malic enzyme (MaeB) and PEP carboxykinase (PckA), as well as fructose 1,6-bisphosphate aldolase (Fba). In contrast, several glycolytic enzymes were upregulated in the Δ*cceR* strain, despite growing on a gluconeogenic carbon substrate. These included a gluconate transporter (GlnT) as well as gluconate kinase (GlnK) and glucokinase (Glk), glucose 6-phosphate isomerase (Pgi), phosphofructokinase (Pfk), and pyruvate kinase (Pyk), as well as three enzymes of the Embden-Meyerhof-Parnas pathway and four enzymes of the Entner-Doudoroff pathway (Zwf, Pgl, Edd, Eda), the main glycolytic route in *P. versutus* (58), a close relative of *P. denitrificans*.

**Figure 7:**
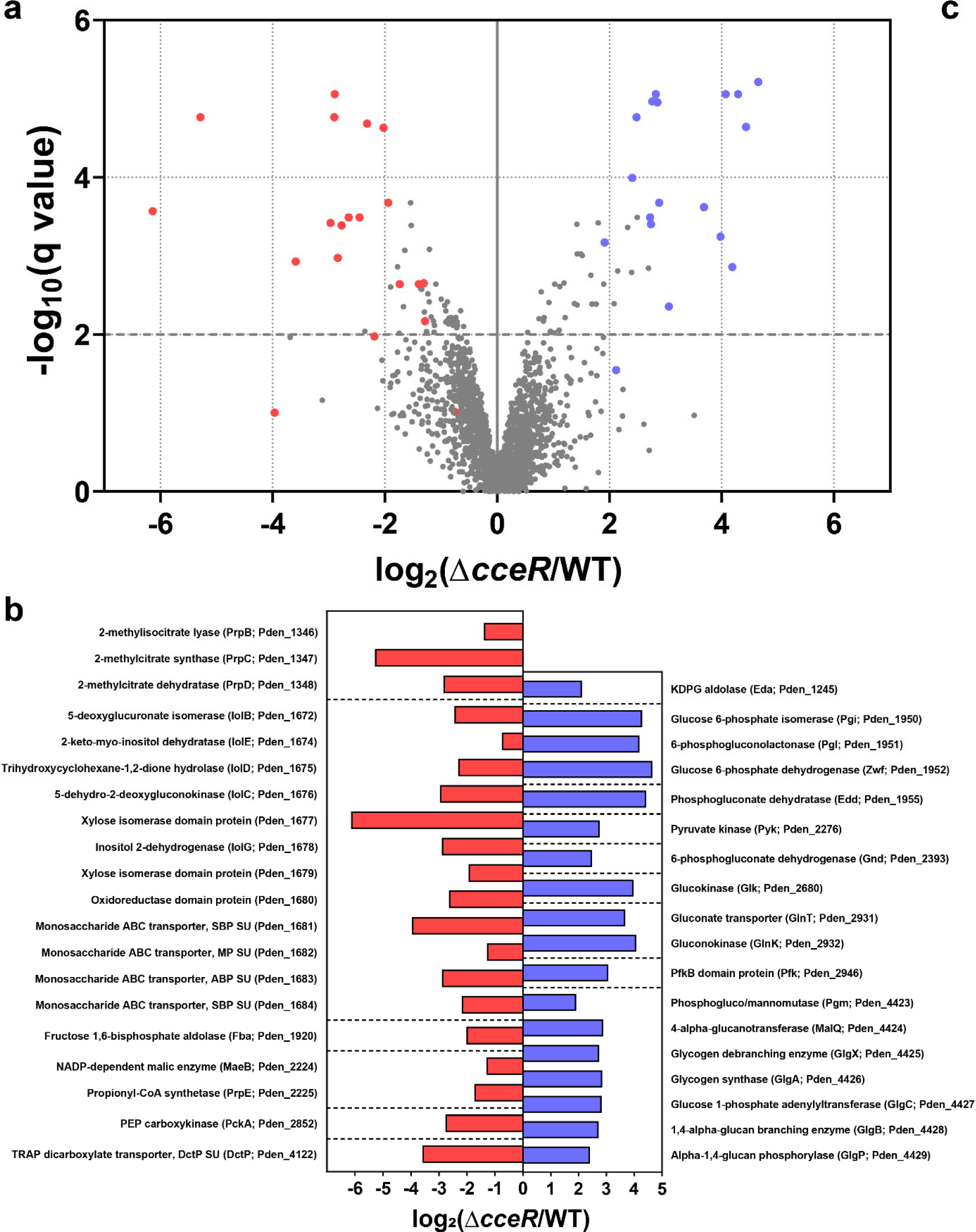
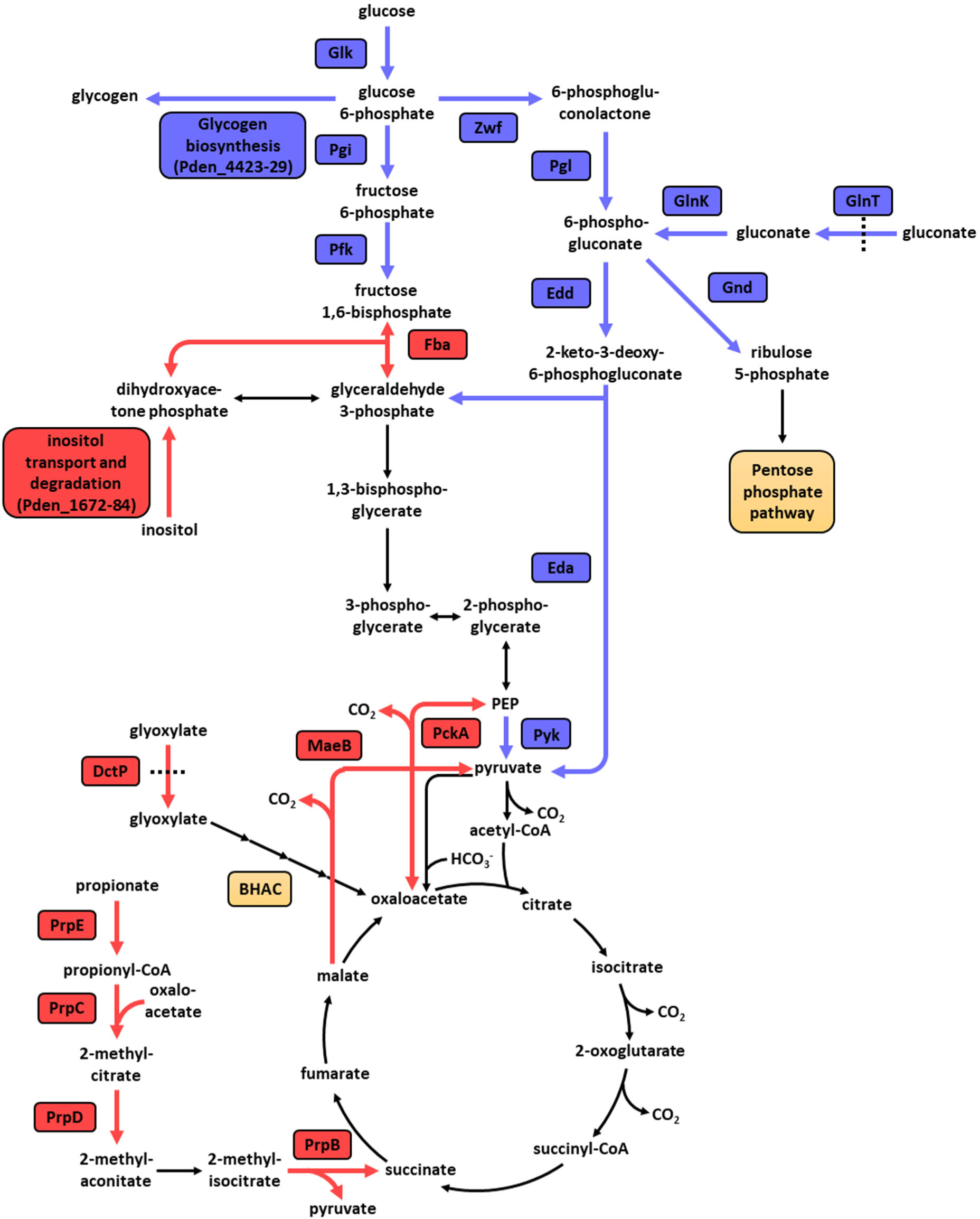
Proteome analysis of *P. denitrificans* DSM 413 Δ*cceR*. **a**, Analysis of the proteome of glyoxylate-grown Δ*cceR* compared to WT. All proteins that were quantified by at least three unique peptides are shown. The proteins in carbon metabolism that showed the strongest decrease or increase in abundance are marked in red or blue in the volcano plot, respectively. X-axis represents log_2_-fold change of the groups means, Y-axis indicates the –log_10_ q value. **b**, The log_2_ fold change of these proteins, sorted by locus name (in brackets). **c**, The role of these up- and downregulated proteins in the carbon metabolism of *P. denitrificans* DSM 413. Altered enzyme production levels in key metabolic routes, such as the Entner-Doudoroff pathway, the C3-C4 node, and the 2-methylcitrate cycle demonstrate marked changes upon deletion of *cceR*.

Based on these results, we concluded that the decreased growth rate of the Δ*cceR* strain on gluconeogenic carbon substrates is due to futile cycling, where glucose is first produced, but then catabolized again by glycolytic enzymes that are constitutively produced in this mutant. In contrast, growth of the Δ*cceR* strain on glycolytic carbon sources is not negatively affected, since high activity of the glycolytic pathways is required for efficient catabolism under these conditions.

Furthermore, our proteomics data supported the conclusion that CceR acts as a repressor of glycolytic pathways and as an activator of gluconeogenic enzymes in *P. denitrificans*, analogous to the role of this regulator in *R. sphaeroides* (57). While the CceR regulon of *P. denitrificans* (determined via proteomics of glyoxylate-grown cultures) is not fully identical to its counterpart in *R. sphaeroides*, there are still large overlaps (**Supplementary Table 2**). Notably, key enzymes in energy metabolism (ATP synthase and NADH dehydrogenase) and the TCA cycle (succinate dehydrogenase, 2-oxoglutarate dehydrogenase, fumarase) are part of the CceR regulon in *R. sphaeroides*, but not in *P. denitrificans*, suggesting that energy conservation and oxidation of acetyl-CoA to CO_2_ are under the control of different regulatory mechanisms in the latter.

Finally, we investigated the substrate uptake hierarchy and substrate consumption rates of the WT and Δ*cceR* strains during growth on glycolytic and gluconeogenic carbon substrates. To this end, these strains were grown on glycolate, glucose, or mixtures thereof, and substrate uptake rates were quantified via LC-MS measurements and luminescence-based assays, respectively.

On glycolate, the Δ*cceR* strain showed a slightly decreased growth rate, as observed before. When growing on glycolate and glucose, uptake of glycolate started and finished earlier than uptake of glucose (**Figure 8a+b**). However, uptake of the two different carbon substrates largely overlapped. Once glycolate was fully consumed, we observed a slightly slower growth phase during which the remaining glucose was used up. This simultaneous uptake of glycolate and glucose was observed for both the WT and Δ*cceR* strains and was independent of initial substrate concentrations. On glycolate and glucose as simultaneous growth substrates, the Δ*cceR* strain was growing similar to the WT strain, which could be explained by the fact that the constitutive activity of glycolytic enzymes was not futile anymore under these conditions. Yet, in all cases, the substrate consumption rates of both glycolate and/or glucose during exponential phase were not significantly changed compared to the WT (**Figure 8c+d**). Overall, this data suggested that CceR controls the glycolysis-gluconeogenesis switch only at the level of the respective assimilation pathways, but not via changes in substrate uptake rate or hierarchy.

**Figure 8:**
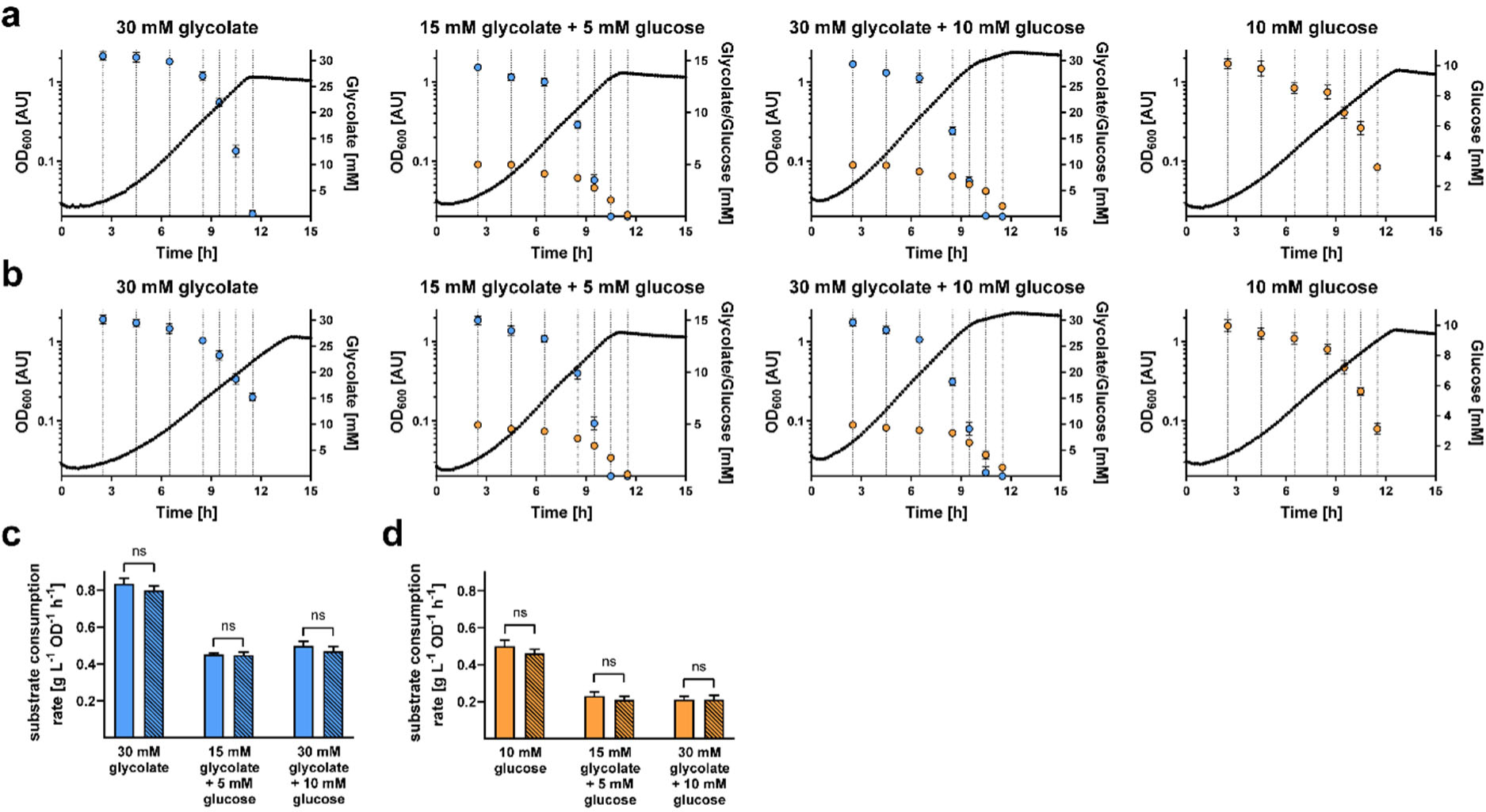
Substrate uptake during growth of *P. denitrificans* DSM 413 WT and Δ*cceR*. The WT (**a**) and Δ*cceR* (**b**) strains were grown on glycolate, glucose, or mixtures of the two carbon sources. At seven time points during growth, glycolate concentrations (light blue) were determined via LC-MS and glucose concentrations (orange) were determined via a luminescence-based assay. The results of *n* = 6 independent experiments are shown; the dot represents the mean, and the error bars represent the standard deviation. Biomass-specific substrate uptake rates were calculated for glycolate (**c**) and glucose (**d**). Empty bars denote the WT strain, striped bars denote the Δ*cceR* strain.

## Discussion

Glycolate and its metabolite glyoxylate are abundant in the environment and are thus readily available carbon sources for heterotrophic microorganisms. Our work aimed at deciphering the regulation of glycolate and glyoxylate assimilation in *P. denitrificans*, an Alphaproteobacterium that relies on glycolate oxidase and the BHAC to funnel these C2 compounds into central carbon metabolism. We determined that BhcR, an IclR-type regulatory protein, controls the BHAC. BhcR is closely related to other glyoxylate-binding regulators and acts as an activator of the *bhc* gene cluster. Furthermore, we discovered that GlcR, a previously unknown member of the GntR family of transcriptional regulators, acts as a repressor to control production of glycolate oxidase. We subsequently extended our investigation towards the regulation of central carbon metabolism in *P. denitrificans* and determined that different carbon substrates are assimilated largely simultaneously, and that the global regulator CceR controls the switch between glycolysis and gluconeogenesis. Taken together, our work elucidates the multi-layered regulatory mechanisms that control assimilation of glycolate and glyoxylate by *P. denitrificans*.

The assimilation of multiple carbon substrates by bacteria has been studied since the seminal work of Monod in the 1940s (59, 60). Bacteria can either consume two nutrients simultaneously or sequentially. Sequential consumption results in a growth curve with two consecutive exponential phases, referred to as diauxie. Both diauxie and simultaneous utilization of two carbon sources are common in microorganisms. The regulatory mechanism responsible for diauxie, known as catabolite repression, allows bacteria to selectively express enzymes for the preferred carbon source even when another one is present (61). The observed simultaneous assimilation of a glycolytic and a gluconeogenic carbon source by *P. denitrificans* can be rationalized based on the conserved topology of central carbon metabolism. When both types of carbon source are present, some precursor molecules for biomass (e.g., glucose 6-phosphate and ribose 5-phosphate) can be synthesized more efficiently from the glycolytic substrate, while other biomass precursors (e.g., oxaloacetate and 2-oxoglutarate) can be synthesized more efficiently from the gluconeogenic substrate. Therefore, it is advantageous for the bacterium to make use of both carbon sources simultaneously (62). Notably, a general growth-rate composition formula that was validated for the growth of *E. coli* on co-utilized glycolytic and gluconeogenic carbon substrates (63) does not seem to be valid for *P. denitrificans* (**Supplementary Table 3**). This might be due to the fact that this formula only takes the regulatory effect of the cAMP-Crp system (64) on catabolic pathways into account. However, the cAMP-Crp system that controls the hierarchical use of different carbon sources is not present in *P. denitrificans*. Therefore, a specific growth-rate composition formula would have to be developed and validated separately for *P. denitrificans* and presumably other Alphaproteobacteria, taking into account the differences in the global regulatory systems that result from the different lifestyles and ecological niches of these versatile microorganisms.

Future work should focus on translating the newly gained knowledge about the transcription factors BhcR and GlcR into the development of robust biosensors for glyoxylate and glycolate, respectively. Established methods for the engineering of sensor modules with a reliable output and applicability for high-throughput screening methods are available (65). A biosensor for the rapid quantification of glycolate would not only be relevant to screen the flux from the CBB cycle into photorespiratory pathways under different conditions, but also to monitor the glycolate output of the CETCH cycle, a promising synthetic pathway for CO_2_ fixation (66, 67).

In summary, our results provide new insights into the regulation of carbon metabolism in *P. denitrificans* and pave the way towards a systems-level understanding of the organism in the future, especially in concert with genome-scale metabolic models that are now available for this bacterium (68, 69).

## Supporting information

Proteomics raw data

## Acknowledgements

We gratefully acknowledge the expert support of Peter Claus in performing small molecule mass spectrometry measurements and Jörg Kahnt in performing mass spectrometry measurements for proteomics.

This study was funded by the Max-Planck-Society (Tobias J. Erb) and the German Research Foundation (SFB987 ‘Microbial diversity in environmental signal response’).

## Competing interests

The authors declare no competing interests.

## Author contributions

L.S.v.B., E.B., and T.J.E. conceptualized the project and designed and supervised experiments. L.S.v.B. performed genetic and biochemical experiments, growth assays, substrate uptake experiments, phylogenetic analysis, and analyzed data. L.H. performed electrophoretic mobility shift assays. K.K. generated and characterized promoter reporter strains. S.B. generated MBP-GlcR and performed fluorescence polarization assays. B.P. generated *P. denitrificans* gene deletion strains. N.P. performed small molecule mass spectrometry. T.G. performed mass spectrometry for proteomics. L.S.v.B. wrote the manuscript, with contributions from all other authors.

## Data availability

All relevant data are available in this article and its Supplementary Information files.

## Notes

### Competing Interest Statement

The authors have declared no competing interest.

