## Supplementary Information for "Multiple levels of transcriptional regulation control glycolate metabolism in *Paracoccus denitrificans*"

*[1]Department of Biochemistry & Synthetic Metabolism, Max Planck Institute for Terrestrial Microbiology, Karl-von-Frisch-Str. 10, D-35043 Marburg, Germany; [2]Institute of Biology Leiden, Leiden University, Sylviusweg 72, 2333 BE Leiden, The Netherlands; [3]Laboratory for Microbiology, Department of Biology, Philipps-University Marburg, Karl-von-Frisch-Str. 8, D-35043 Marburg, Germany; [4]Facility for Mass Spectrometry and Proteomics, Max Planck Institute for Terrestrial Microbiology, Karl-von-Frisch-Str. 10, D-35043 Marburg, Germany; [5]Facility for Metabolomics and Small Molecule Mass Spectrometry, Max Planck Institute for Terrestrial Microbiology, Karl-von-Frisch-Str. 10, 35043 Marburg, Germany; [6]LOEWE-Center for Synthetic Microbiology, Philipps-University Marburg, Karl-von-Frisch-Str. 8, D-35043 Marburg, Germany.*

**Table of Contents Supplementary Information:**

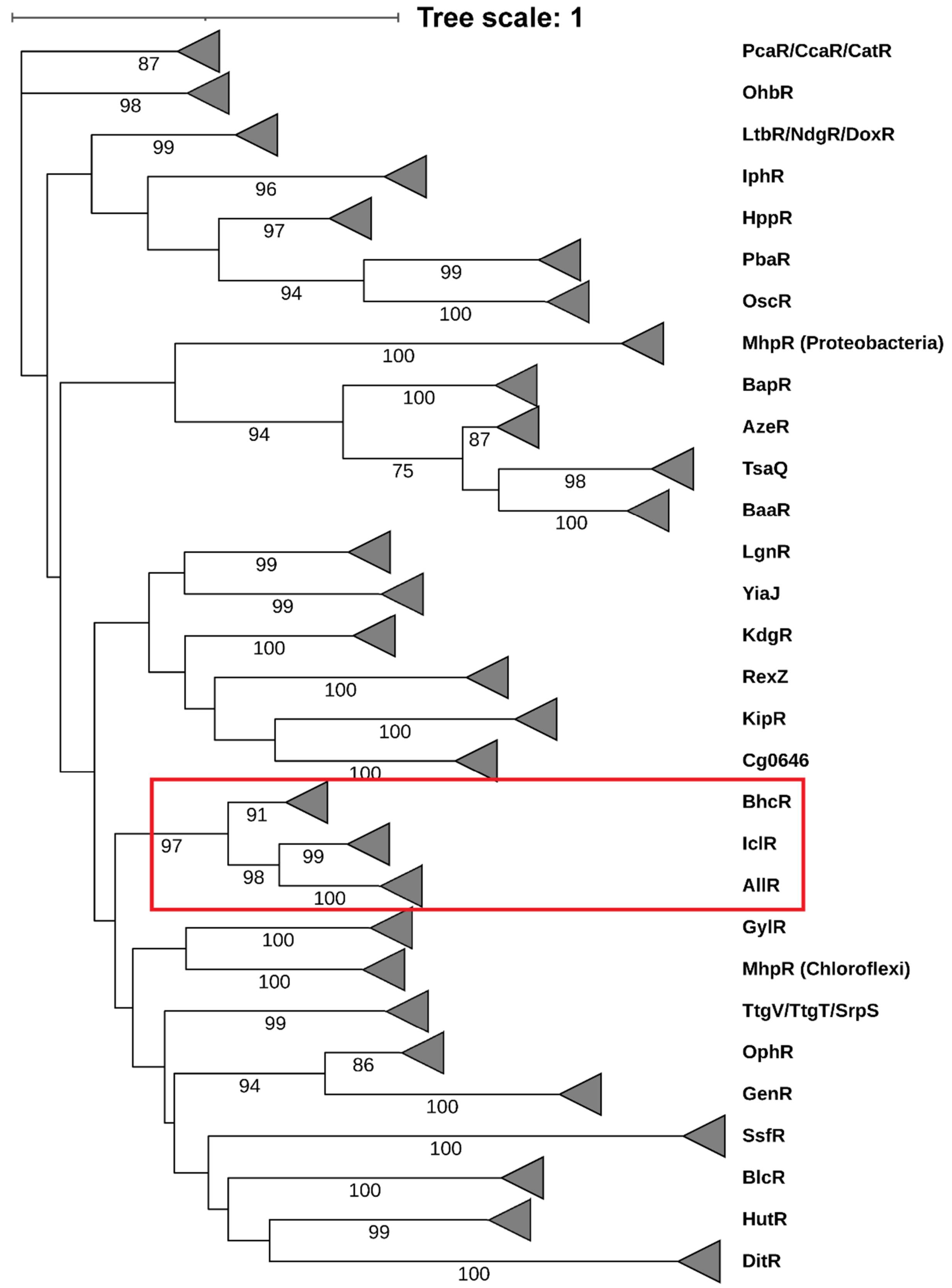

**Supplementary Figure 1: Maximum likelihood phylogenetic tree of the IclR family of transcriptional regulators.** Sequences of the transcription factor BhcR and its homologs form a distinct clade within the branch of glyoxylate-binding transcription factors (denoted by a red frame), which also includes the AllR and IclR subfamilies. Bootstrap values of at least 50 are given on the respective nodes.

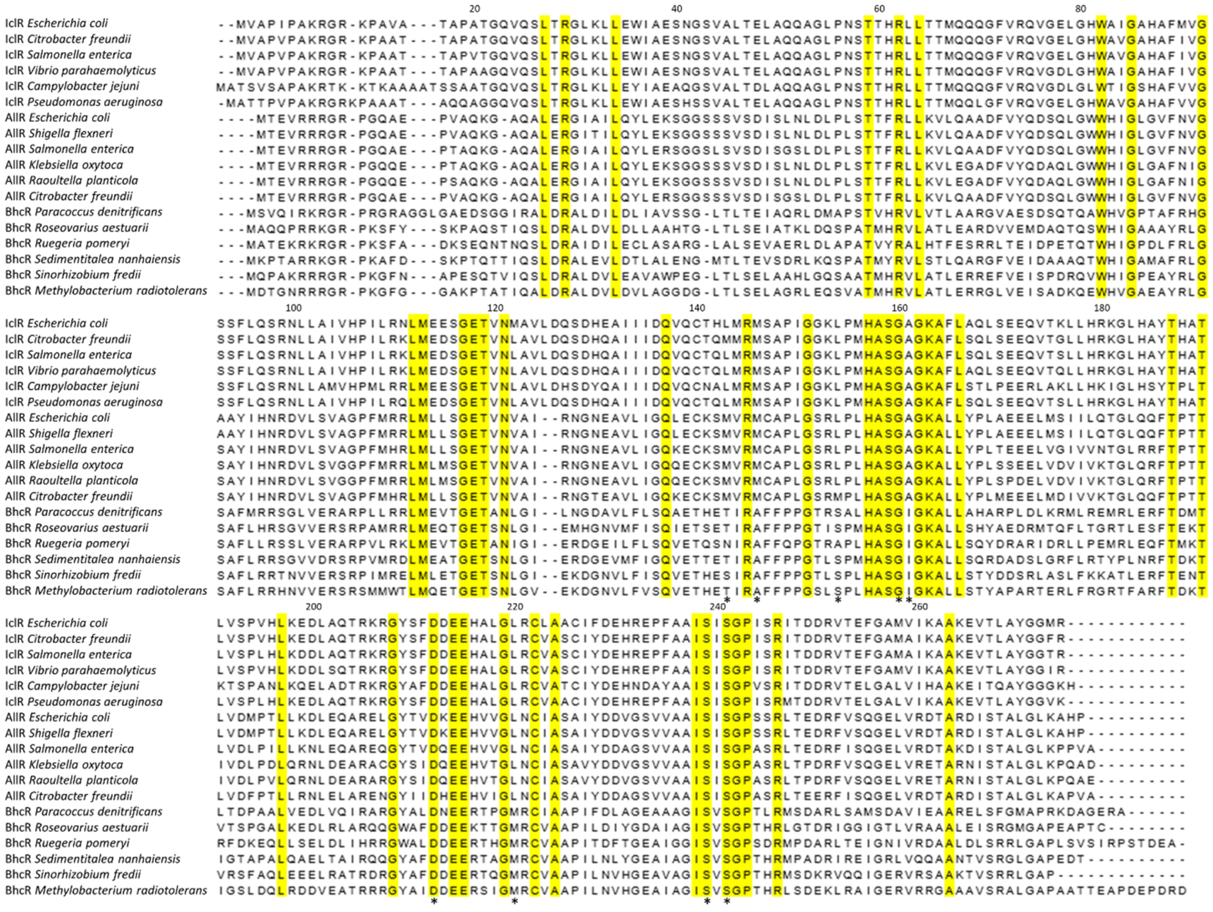

**Supplementary Figure 2: Alignment of IclR, AllR, and BhcR amino acid sequences.** Amino acids that are fully conserved across this dataset are highlighted in yellow. Amino acids that were previously identified as ligand-binding residues in IclR are denoted with asterisks below the alignment. Numbering of amino acids above the alignment is based on the sequence of *E. coli* IclR.

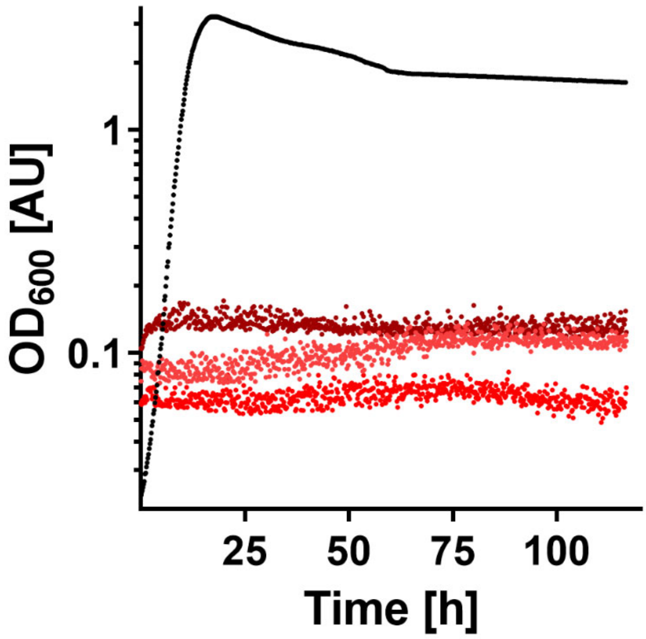

**Supplementary Figure 3:** ***Paracoccus denitrificans*** **is not capable of growth on oxalate.** No growth was observed on 5 mM (light red), 10 mM (red), or 30 mM oxalate (dark red) as sole source of carbon and energy. Growth on 40 mM pyruvate (black) was monitored as a positive control.

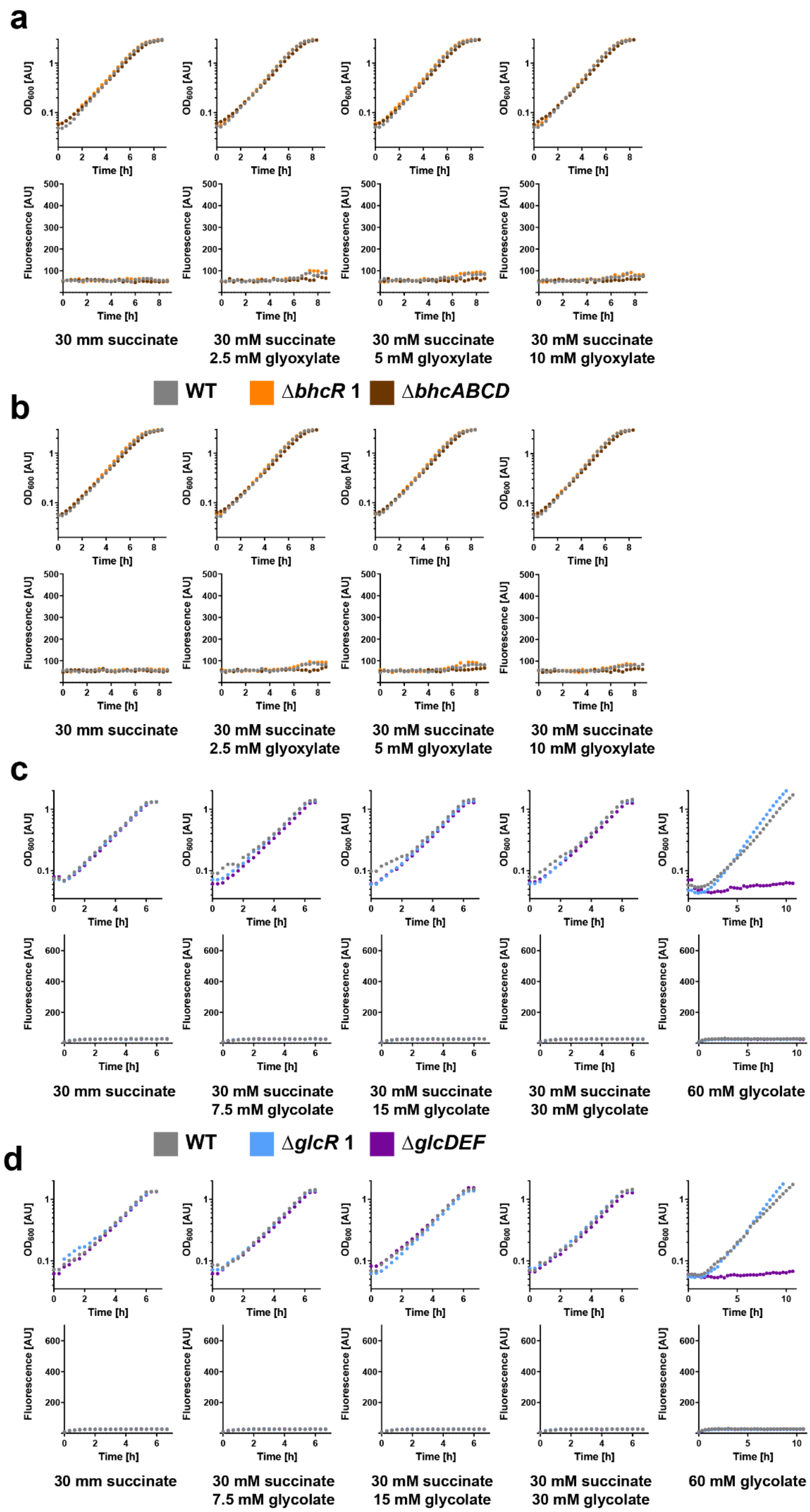
a
OD600 [AU]
Time [h]
Fluorescence [AU]
30 mm succinate
30 mM succinate
2.5 mM glyoxylate
30 mM succinate
5 mM glyoxylate
30 mM succinate
10 mM glyoxylate
WT
ΔbhcR 1
ΔbhcABCD
b
30 mm succinate
30 mM succinate
2.5 mM glyoxylate
30 mM succinate
5 mM glyoxylate
30 mM succinate
10 mM glyoxylate
c
30 mm succinate
30 mM succinate
7.5 mM glycolate
30 mM succinate
15 mM glycolate
30 mM succinate
30 mM glycolate
60 mM glycolate
WT
ΔglcR 1
ΔglcDEF
d
30 mm succinate
30 mM succinate
7.5 mM glycolate
30 mM succinate
15 mM glycolate
30 mM succinate
30 mM glycolate
60 mM glycolate

**Supplementary Figure 4: Characterization of *P. denitrificans* promoter reporter strains (negative controls). a, b**, Growth and fluorescence of promoter reporter strains Δ*bhcR* (orange), Δ*bhcABCD* (brown), and WT (grey) without a plasmid (**a**) or with pTE714 (**b**) on different carbon sources. **c, d**, Growth and fluorescence of promoter reporter strains Δ*glcR* (light blue), Δ*glcDEF* (purple), and WT (grey) without a plasmid (**c**) or with pTE714 (**d**) on different carbon sources. All experiments were repeated three times independently with similar results.

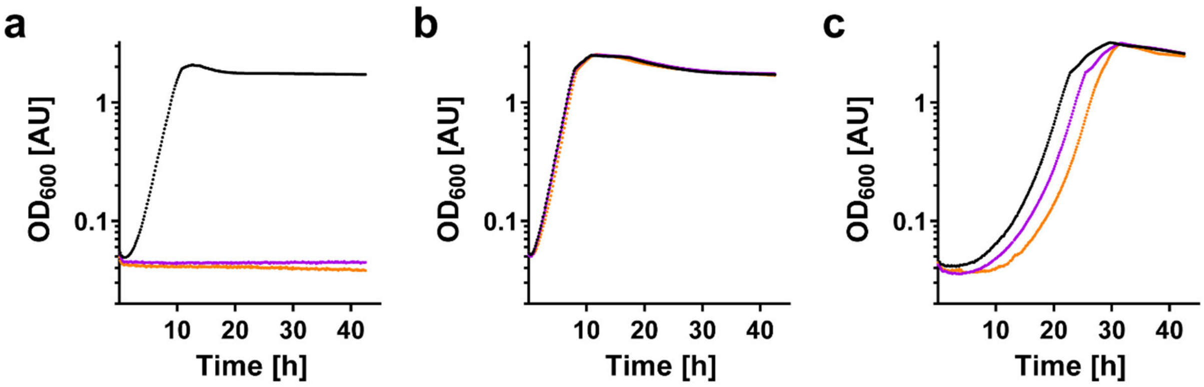

**Supplementary Figure 5: Growth of *P. denitrificans* Δ*glcDEF*.** Two Δ*glcDEF* deletion strains, in which the genes Pden_4397-99 were replaced with a kanamycin resistance cassette in either the same or the opposite direction of transcription (pink, orange) were unable to grow on 60 mM glycolate (**a**). In contrast, these two deletion strains grew similarly to the WT strain (black) on 30 mM succinate (**b**) and 60 mM acetate (**c**).

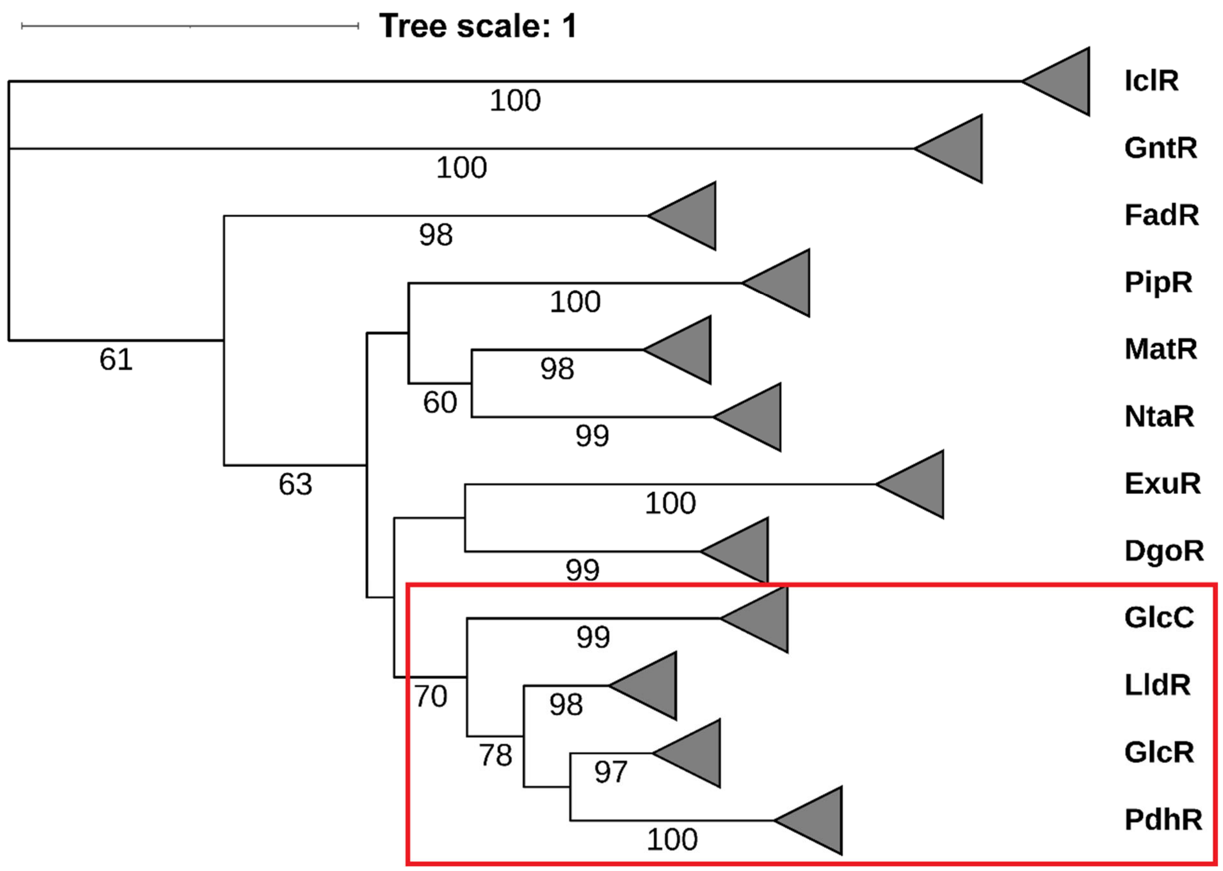

**Supplementary Figure 6: Maximum likelihood phylogenetic tree of the FadR subfamily of transcriptional regulators.** Sequences of IclR-type and GntR-type regulators were used as outgroups. Sequences of the regulator GlcR and its homologs form a distinct clade within the branch of transcription factors that interact with glycolate, pyruvate, or lactate (denoted by a red frame), which also includes the GlcC, LldR, and PdhR subfamilies. Bootstrap values of at least 50 are given on the respective nodes.

**Supplementary Table 1: The *glcRDEF* gene cluster in various alphaproteobacterial strains.** Gene IDs for the respective *glcRDEF* genes are according to the respective genome sequences deposited in the NCBI Nucleotide database (https://www.ncbi.nlm.nih.gov/nucleotide/).

| Strain | Order | *glcR* | *glcD* | *glcE* | *glcF* |
|---|---|---|---|---|---|
| *Paracoccus denitrificans* DSM 413 | *Rhodobacterales* | Pden_4400 | Pden_4399 | Pden_4398 | Pden_4397 |
| *Paracoccus pantotrophus* DSM 2944 | *Rhodobacterales* | ESD82_RS02595 | ESD82_RS02600 | ESD82_RS02605 | ESD82_RS02610 |
| *Paracoccus zeaxanthinifaciens* ATCC 21588 | *Rhodobacterales* | F804_RS0111220 | F804_RS0111210 | F804_RS0111205 | F804_RS0111200 |
| *Paracoccus methylovorus* H4-D09 | *Rhodobacterales* | JWJ88_RS16105 | JWJ88_RS16110 | JWJ88_RS16115 | JWJ88_RS16120 |
| *Methylarcula marina* VKM B-2159 | *Rhodobacterales* | A7A09_RS08100 | A7A09_RS08105 | A7A09_RS08110 | A7A09_RS08115 |
| *Puniceibacterium antarcticum* SM1211 | *Rhodobacterales* | P775_RS11050 | P775_RS11045 | P775_RS11040 | P775_RS11035 |
| *Rhodobacter xinxiangensis* TJ48 | *Rhodobacterales* | E2K76_RS04305 | E2K76_RS04315 | E2K76_RS04320 | E2K76_RS04325 |
| *Chenggangzhangella methanolivorans* CHL1 | *Rhizobiales* | K6K41_RS14170 | K6K41_RS14165 | K6K41_RS14160 | K6K41_RS14155 |
| *Afipia* sp. 1NLS2 | *Rhizobiales* | AFIDRAFT_RS03560 | AFIDRAFT_RS03555 | AFIDRAFT_RS03550 | AFIDRAFT_RS03545 |

**Supplementary Table 2: The CceR regulon in *P. denitrificans* and *Rhodobacter sphaeroides*.** Data for *R. sphaeroides* and the predicted CceR regulon of *P. denitrificans* were previously published (1).

| protein (gene) | *R. sphaeroides* | *P. denitrificans* (predicted) | *P. denitrificans* (experimentally determined) |
|---|---|---|---|
| PEP carboxykinase (*pckA*) | X | | X |
| malate dehydrogenase (*mdh*) | X | | |
| succinate dehydrogenase (*sdhDA*) | X | | |
| ATP synthase (*atpBEFHAC*) | X | | |
| fructose 1,6-bisphosphatase (*fbp*) | X | | |
| succinyl-CoA synthetase (*sucCD*) | X | | |
| 2-OG dehydrogenase (*sucB*) | X | | |
| fumarase (*fumC*) | X | | |
| NADH dehydrogenase (*nuoL*) | X | | |
| fructose 1,6-bisphosphate aldolase (*fba*) | X | | X |
| pyruvate dehydrogenase (*pdhAB*) | X | | |
| glucose 6-phosphate dehydrogenase (*zwf*) | X | | X |
| 6-phosphogluconolactonase (*pgl*) | X | | X |
| glucose 6-phosphate isomerase (*pgi*) | X | | X |
| phosphogluconate dehydratase (*edd*) | X | X | X |
| KDPG aldolase (*eda*) | X | X | X |
| cytochrome c-554 (*cycF*) | X | | |
| pyruvate kinase (*pyk*) | | X | X |
| TRAP dicarboxylate transporter (*dctP*) | | X | X |
| gluconate transporter (*glnT*) | | X | X |
| gluconokinase (*glnK*) | | X | X |
| alpha-1,4-glucan phosphorylase (*glgP*) | | X | X |
| 1,4-alpha-glucan branching enzyme (*glgB*) | | X | X |
| glucose 1-phosphate adenylyltransferase (*glgC*) | | X | X |
| glycogen synthase (*glgA*) | | X | X |
| glycogen debranching enzyme (*glgX*) | | X | X |
| 4-alpha-glucanotransferase (*malQ*) | | X | X |
| phosphogluco/mannomutase (*pgm*) | | X | X |
| phosphofructokinase (*pfk*) | | | X |
| glucokinase (*glk*) | | | X |
| 6-phosphogluconate dehydrogenase (*gnd*) | | | X |
| propionyl-CoA synthetase (*prpE*) | | | X |
| 2-methylisocitrate lyase (*prpB*) | | | X |
| 2-methylisocitrate synthase (*prpC*) | | | X |
| 2-methylisocitrate dehydratase (*prpD*) | | | X |
| malic enzyme (*maeB*) | | | X |
| inositol degradation pathway (Pden_1672-1684) | | | X |

**Supplementary Table 3: Calculated and experimentally determined growth rates of *P. denitrificans* on minimal medium with glycolytic and gluconeogenic carbon substrates.** The growth rate composition formula used here was previously developed and validated for *E. coli* (2). All experimentally determined growth rates are averages from six independent experiments; variability between independent experiments was not more than 5%. For the calculations, a $\lambda_C$ value of 0.85 was estimated for *P. denitrificans*.

| **Growth rate μ ($h^{-1}$)** | **as sole carbon source** | **with 10 mM glucose (calculated)** | **with 10 mM glucose (experimentally determined)** |
|---|---|---|---|
| **30 mM glycolate** | 0.51 | 0.60 | 0.51 |
| **30 mM glyoxylate** | 0.28 | 0.48 | 0.39 |
| **10 mM glucose** | 0.38 | - | - |

**Supplementary Table 4 | Strains used in this study**

| **strain** | **genotype or relevant features**[a] | **source or reference** |
|---|---|---|
| *E. coli* DH5α | *supE44*, *ΔlacU169* (*Φ80lacZDM15*), *hsdR17*, *recA1*, *endA1*, *gyrA96*, *thi*-1, *relA1* | Thermo Fisher Scientific, Waltham, USA |
| *E. coli* ST18 | *pro*, *thi*, *hsdR1*,$Tp^R$, $Sm^R$; chromosome::RP4-2, Tc::Mu-Kan::Tn7/λpir, λpir, Δ*hemA* | Thoma and Schobert (3) |
| *E. coli* BL21 AI | *ompT*, *gal*, *dcm*, *lon*, $hsdS_B(r_B^- m_B^-)$, $[malB^+]_{K-12}(\lambda^S)$, *araB*::*T7RNAP-tetA* | Thermo Fisher Scientific, Waltham, USA |
| *P. denitrificans* DSM 413 | WT strain | Beijerinck and Minkman (4) |
| *P. denitrificans* DSM 413 Δ*bhcR* 1 | Δ3922; $Km^R$ (orientation 1) | this work |
| *P. denitrificans* DSM 413 Δ*bhcR* 2 | Δ3922; $Km^R$ (orientation 2) | this work |
| *P. denitrificans* DSM 413 Δ*bhcABCD* 1 | Δ3921-18; $Km^R$ (orientation 1) | Schada von Borzyskowski, Severi (5) |
| *P. denitrificans* DSM 413 Δ*bhcABCD* 2 | Δ3921-18; $Km^R$ (orientation 2) | Schada von Borzyskowski, Severi (5) |
| *P. denitrificans* DSM 413 Δ*glcR* 1 | Δ4400; $Km^R$ (orientation 1) | this work |
| *P. denitrificans* DSM 413 Δ*glcR* 2 | Δ4400; $Km^R$ (orientation 2) | this work |
| *P. denitrificans* DSM 413 Δ*glcDEF* 1 | Δ4399-97; $Km^R$ (orientation 1) | this work |
| *P. denitrificans* DSM 413 Δ*glcDEF* 2 | Δ4399-97; $Km^R$ (orientation 2) | this work |
| *P. denitrificans* DSM 413 Δ*cceR* 1 | Δ1978; $Km^R$ (orientation 1) | this work |
| *P. denitrificans* DSM 413 Δ*cceR* 2 | Δ1978; $Km^R$ (orientation 2) | this work |

[a] $Km^R$, kanamycin resistance; $Tp^R$, trimethoprim resistance; $Sm^R$, streptomycin resistance

**Supplementary Table 5 | Plasmids used in this study**

| plasmid | relevant features[a] | source or reference |
|---|---|---|
| pET16b | *E. coli* expression vector, T7 promoter, $Amp^R$ | Merck Chemicals GmbH, Darmstadt, Germany |
| pET16b-BhcR | expression vector for N-terminally His-tagged BhcR, $Amp^R$ | Schada von Borzyskowski, Severi (5) |
| pET16b-GlcR | expression vector for N-terminally His-tagged GlcR, $Amp^R$ | this work |
| pMBP-sfgfp_dropout (pTE5400) | expression vector for N-terminally His-tagged-maltose-binding protein (10x-His-MBP) with sfgfp dropout, compatible with Marburg Collection (6), $Cam^R$ | this work |
| pMBP-GlcR (pTE5418) | expression vector for N-terminally His- and MBP-tagged GlcR, $Cam^R$ | this work |
| pREDSIX | mobilizable, high-copy-number cloning and mutagenesis vector; $Amp^R$ | Ledermann, Strebel (7) |
| pRGD-KmR | donor vector for resistance gene, *aphII* in polylinker; $Amp^R$, $Km^R$ | Ledermann, Strebel (7) |
| pREDSIX-*bhcR-1* | knockout vector for the *bhcR* gene in *P. denitrificans* DSM 413; $Km^R$ (orientation 1) | this work |
| pREDSIX-*bhcR-2* | knockout vector for the *bhcR* gene in *P. denitrificans* DSM 413; $Km^R$ (orientation 2) | this work |
| pREDSIX-gl*cR-1* | knockout vector for the *glcR* gene in *P. denitrificans* DSM 413; $Km^R$ (orientation 1) | this work |
| pREDSIX-gl*cR-2* | knockout vector for the *glcR* gene in *P. denitrificans* DSM 413; $Km^R$ (orientation 2) | this work |
| pREDSIX-*cceR-1* | knockout vector for the *cceR* gene in *P. denitrificans* DSM 413; $Km^R$ (orientation 1) | this work |
| pREDSIX-*cceR-2* | knockout vector for the *cceR* gene in *P. denitrificans* DSM 413; $Km^R$ (orientation 2) | this work |
| pREDSIX-*glcDEF-1* | knockout vector for the *glcDEF* gene cluster in *P. denitrificans* DSM 413; $Km^R$ (orientation 1) | this work |
| pREDSIX-*glcDEF-2* | knockout vector for the *glcDEF* gene cluster in *P. denitrificans* DSM 413; $Km^R$ (orientation 2) | this work |
| pTE100 | empty expression vector for Alphaproteobacteria; $Tc^R$ | Schada von Borzyskowski, Remus-Emsermann (8) |
| pTE714 | promoter probe vector for Alphaproteobacteria based on pTE100 containing a RBS and mCherry; $Tc^R$ | this work |
| pTE714_3922/3921_ig | promoter probe vector containing mCherry under control of the promoter located in the intergenic region between Pden_3922 and Pden_3921; $Tc^R$ | this work |
| pTE714_4400/4399_ig | promoter probe vector containing mCherry under control of the promoter located in the intergenic region between Pden_4400 and Pden_4399; $Tc^R$ | this work |

[a] $Km^R$, kanamycin resistance; $Amp^R$, ampicillin resistance; $Tc^R$, tetracycline resistance; $Cam^R$, chloramphenicol resistance

### Supplementary Table 6 | Primers used in this study

| target | name | sequence[a] | cut site |
|---|---|---|---|
| Pden_*glcR* | glcR_16b_fw | 5'-GACGCTG**CATATG**ACCAACGCATCCGTGAAATCC-3' | *Nde*I |
| Pden_*glcR* | glcR_16b_rv | 5'-GACACTC**GGATCC**CTAGCCCTCTGCGGCCGGGGTGG-3' | *Bam*HI |
| Pden_*bhcR*_up | bhcR_up_fw | 5'-GGTCTGACAGGTTTAAACTCTAGACCGGAAATCCATCGAACCGATG-3' | --- |
| Pden_*bhcR*_up | bhcR_up_rv | 5'-AAGTTTAGACGAA**GGTACC**TCAATTTTTCCTTTCGACAAC-3' | *Kpn*I |
| Pden_*bhcR*_down | bhcR_down_fw | 5'-AAATTGA**GGTACC**TTCGTCTAAACTTGACCAGGACATGCCC-3' | *Kpn*I |
| Pden_*bhcR*_down | bhcR_down_rv | 5'-CTTAAGGCTAGCATGCATCCTAGGCGGGCTGTAGCCGGGCCCGATC-3' | --- |
| Pden_*glcR*_up | glcR_up_fw | 5'-GGTCTGACAGGTTTAAACTCTAGACAGGCCCCATCTCGACCCCAC-3' | --- |
| Pden_*glcR*_up | glcR_up_rv | 5'-GGCCAGTCCTCT**CATATG**GGGCGCTCATGCGGTTGTCG-3' | *Nde*I |
| Pden_*glcR*_down | glcR_down_fw | 5'-AGCGCCC**CATATG**AGAGGACTGGCCGGCAAGGAAAAG-3' | *Nde*I |
| Pden_*glcR*_down | glcR_down_rv | 5'-CTTAAGGCTAGCATGCATCCTAGGCGATGGCGACCGTCGCCACCC-3' | --- |
| Pden_*cceR*_up | cceR_up_fw | 5'-GGTCTGACAGGTTTAAACTCTAGACTCCGCAAGGCCGGCCAGGAC-3' | --- |
| Pden_*cceR*_up | cceR_up_rv | 5'-GCAGGCGCGGCG**CATATG**CGCAACTACCGGGACGC-3' | *Nde*I |
| Pden_*cceR*_down | cceR_down_fw | 5'-AGTTGCG**CATATG**CGCCGCGCCTGCGGGCGGCG-3' | *Nde*I |
| Pden_*cceR*_down | cceR_down_rv | 5'-CTTAAGGCTAGCATGCATCCTAGGCCATGCCAGCTGCATCAGCGTGA ACCATGTCGTCTG-3' | --- |
| Pden_*glcDEF*_up | glcDEF_up_fw | 5'-GGTCTGACAGGTTTAAACTCTAGACCCGCATGACCAACGCCACCGTG-3' | --- |
| Pden_*glcDEF*_up | glcDEF_up_rv | 5'-AAGGGAGGAGAGC**CATATG**GCTCGCCCCGGCTTGGGAACG-3' | *Nde*I |
| Pden_*glcDEF*_down | glcDEF_down_fw | 5'-AGCCGGGGCGAGC**CATATG**GCTCTCCTCCCTTGGTCGAACGG-3' | *Nde*I |
| Pden_*glcDEF*_down | glcDEF_down_rv | 5'-CTTAAGGCTAGCATGCATCCTAGGCCTGGCGCAGCAACTCGCGGAT-3' | --- |
| mCherry | mCherry_fw | 5'-GACACGC**CATATG**GTGAGCAAG-3' | *Nde*I |
| mCherry | mCherry_rv | 5'-GCTACTC**GAATTC**TTACTTGTACAGCTCGTCCATGC-3' | *Eco*RI |
| Pden_3922/3921_ig | Pden3922_ig_rv | 5'-GATAT**GAATTC**TCAGCCGCCGGCACGTCCC-3' | *Eco*RI |
| Pden_3922/3921_ig | Pden3921_ig_rv | 5'-CG**TCTAGA**TCACTCGGGGATGTTGGTCGG-3' | *Xba*I |
| Pden_4400/4399_ig | Pden4400_ig_rv | 5'-GATAT**GAATTC**TCACTGGGCCACGGCCTCG-3' | *Eco*RI |
| Pden_4400/4399_ig | Pden4399_ig_rv | 5'-CTT**TCTAGA**TCACGCACGCGCCAGGATGCC-3' | *Xba*I |
| Pden_*Pbhc* | Pbhc_fw | 5'-CCCTTTTGCGGATTTGAACCG-3' | --- |
| Pden_*Pbhc* | Pbhc_rev-dye[b] | 5'-GGAATGAAGATCGGGTTCTGGC-3' | --- |
| Pden_*bhcA* | bhcA_fw | 5'-GGGGTCAAATCGGACATCGC-3' | --- |
| Pden_*bhcA* | bhcA_rev-dye[b] | 5'-GCGGATGTCGAAGAAGGTGC-3' | --- |
| Pden_*Pglc* | Pglc_fw | 5'-CAAGCTCTCCTCCCTTGGTCGAACG-3' | --- |
| Pden_*Pglc* | Pglc_rev-dye[b] | 5'-CATGGGCGCTCATGCGGTTGTC-3' | --- |
| Pden_*glcD* | glcD_fw | 5'-GCGATAGGCGGTCAGCGCGTCGCATTCATAG-3' | --- |
| Pden_*glcD* | glcD_rev-dye[b] | 5'-TTGTCCGGTATTGCCATGCC-3' | --- |
| Pden_*Pglc*[c] | [6FAM]-Pglc_fw | 5'-[6FAM]-GGGCGCTCATGCGGTTGTC-3' | --- |
| Pden_*Pglc*[c] | Pglc_rev | 5'-GCTCTCCTCCCTTGGTCGAAC-3' | --- |
| -[c] | [6FAM]-tetO_fw | 5'-[6FAM]-TCCCTATCAGTGATAGAGA-3' | --- |
| -[c] | tetO_rev | 5'-TCTCTATCACTGATAGGGA-3' | --- |

[a] Nucleotides in bold and underlined are recognition sites for endonuclease restriction enzymes. [b] 5'-labeled with Dyomics 781 fluorescent dye. [c] used to generate fluorescence polarization templates.
